# mTORC1 IN THE ORBITOFRONTAL CORTEX PROMOTES HABITUAL ALCOHOL SEEKING

**DOI:** 10.1101/564310

**Authors:** Nadege Morisot, Khanhky Phamluong, Yann Ehinger, Anthony L. Berger, Jeffrey J. Moffat, Dorit Ron

## Abstract

The mechanistic target of rapamycin complex 1 (mTORC1) plays an important role in dendritic translation and in learning and memory. We previously showed that heavy alcohol use activates mTORC1 in the orbitofrontal cortex (OFC) of rodents (1). Here, we set out to determine the consequences of alcohol-dependent mTORC1 activation in the OFC. We found that inhibition of mTORC1 activity attenuates alcohol seeking and restores sensitivity to outcome devaluation in rats that habitually seek alcohol. In contrast, habitual responding for sucrose was unaltered by mTORC1 inhibition, suggesting that mTORC1’s role in habitual behavior is specific to alcohol. We further show that inhibition of GluN2B in the OFC attenuates alcohol-dependent mTORC1 activation, alcohol seeking and habitual responding for alcohol. Together, these data suggest that the GluN2B/mTORC1 axis in the OFC drives alcohol seeking and habit.

## INTRODUCTION

mTORC1 is a multiprotein complex that contains the serine/threonine protein kinase mTOR, the regulatory associated protein of TOR (Raptor), and other enzymes and adaptor proteins (2). Upon activation, mTORC1 phosphorylates eIF4E-binding protein (4E-BP) and the ribosomal protein S6 kinase (S6K), which in turn phosphorylates its substrate, S6 (3). These phosphorylation events lead to the translation of a subset of mRNAs to proteins (2, 3). In the CNS, mTORC1 is responsible for the local dendritic translation of synaptic proteins (4, 5). As such, mTORC1 plays an important role in synaptic plasticity, and learning and memory (6).

Increasing lines of evidence in rodents suggest that mTORC1 is a key player in mechanisms underlying alcohol use disorder (AUD) (7). For instance, excessive alcohol intake and reinstatement of alcohol place preference activate mTORC1 in the rodent nucleus accumbens (NAc) (1, 8–10), resulting in the translation of synaptic proteins, which in turn produce synaptic and structural adaptations that drive and maintain heavy alcohol use and relapse (1, 8, 9, 11, 12). Repeated cycles of voluntary binge drinking of alcohol and withdrawal also produces a robust and sustained activation of mTORC1 in the OFC of rodents (1); however, the behavioral consequences of this activation are unknown.

The OFC integrates sensory and reward information (13) and is an essential player in decision making (13–15), in stimulus-outcome association (16–18), in updating the value of predicted outcomes (15, 19), in goal-directed behaviors (16, 20, 21), and in compulsive behavior (22, 23). Exposure to drugs of abuse impairs the performance of OFC-dependent behavioral tasks (24), and aberrant neuroadaptations induced by drugs of abuse in the OFC contribute to drug-seeking and compulsive drug taking (24–26). For example, withdrawal from alcohol vapor exposure produces morphological and cellular alterations in rodents (27, 28). Inactivation of the OFC enhances alcohol consumption in alcohol-dependent mice (29), and chronic alcohol use alters the protein landscape in the primate OFC (30). In humans, degraded OFC white matter, and reduced neuronal density are associated with alcohol-dependence (31, 32), and presentation of alcohol-associated cues to alcohol-dependent subjects elicits OFC activation (33–35).

Here, using rats as a model system, we set out to examine whether mTORC1 in the OFC contributes to the mechanisms underlying AUD.

## METHODS

### Reagents

Anti-phospho-S6 (S235/236, 1:500), anti-S6 (1:1000), anti-phospho-4E-BP (T37/46, 1:500), anti-4E-BP (1:1000), anti-phospho-CaMKII (T286, 1:1000), anti-phospho-S6K (T389, 1:500) and anti-S6K (1:500) antibodies were purchased from Cell Signaling Technology (Danvers, MA). Anti-CaMKII (1:500) antibodies were purchased from Santa Cruz Biotechnology (Santa Cruz, CA). Anti-Actin (1:10,000) antibodies, phosphatase Inhibitor Cocktails 2 and 3, Dimethyl sulfoxide (DMSO), D-sucrose and NMDA were from Sigma Aldrich (St. Louis, MO). Nitrocellulose membranes and anti-NeuN antibodies were purchased from EMD Millipore (Billerica, MA, USA). Enhanced Chemiluminescence (ECL) was purchased from GE Healthcare (Pittsburg, PA, USA). Donkey anti-rabbit horseradish peroxidase (HRP), and donkey anti-mouse horseradish peroxidase (HRP) were purchased from Jackson ImmunoResearch (West Grove, PA). EDTA-free complete mini Protease Inhibitor Cocktails were purchased from Roche (Indianapolis, IN). NuPAGE Bis-Tris precast gels and Phosphate buffered saline (PBS) were purchased from Life Technologies (Grand Island, NY). Pierce Bicinchoninic Acid (BCA) protein assay kit was obtained from Thermo Scientific (Rockford, IL), and ProSignal Blotting Film were purchased from Genesee Scientific (El Cajon, CA). Ethyl alcohol (190 proof) was purchased from VWR (Radnor, PA). Rapamycin was purchased from LC laboratories (Woburn, MA). Ro25-6981 was purchased from Tocris Bioscience (Bristol, UK).

### Subjects

Male Long Evans rats (Harlan, Indianapolis, IN) were one (*ex vivo* experiment) or two months (*in vivo* experiments) old at their arrival. Animals were single-housed in a temperature- and humidity-controlled colony room (22±2 °C, relative humidity: 50–60%) under a normal 12h light/dark cycle (lights on at 7:00AM) with food and water available ad libitum. Rats were given a week of habituation to the housing conditions before the beginning of the experiments. All animal procedures were approved by the University of California San Francisco Institutional Animal Care and Use Committee (IACUC) and were conducted in agreement with the Association for Assessment and Accreditation of Laboratory Animal Care (AAALAC).

### Preparation of solutions

Alcohol was diluted to 20% (v/v) and sucrose to 1-8% (w/v) in tap water. Rapamycin (50ng/μl) was dissolved in PBS. Ro25-6981 (10μg/μl) and NMDA (25μM) were dissolved in 0.1% DMSO in PBS.

### Intermittent access to 20% alcohol using two bottle choice and drug infusion

Rats underwent intermittent access to 20% alcohol in a 2-bottle choice (IA-20%2BC) paradigm for 7 weeks as described in (36). Specifically, rats were given 24 hours of concurrent access to one bottle of 20% alcohol (v/v) in tap water and one bottle of water. Control rats had access to water only. Drinking sessions started at 12:00pm on Monday, Wednesday and Friday, with 24- or 48 hours (weekend) of alcohol-deprivation periods in which rats consumed only water. The placement (left or right) of the water or alcohol solution was alternated between each session to control for side preference. Water and alcohol bottles were weighed at the beginning and at the end of each alcohol drinking session. Rats were weighed once a week. Rats drinking more than 3.5 g/kg/24h on the last week of IA-20%2BC were selected for the study, approximately 70-80% of the cohort.

### Stereotaxic Surgery

Following stable baseline of alcohol drinking, rats underwent stereotaxic surgery. Specifically, rats were continuously anesthetized using isoflurane (Baxter) and bilateral guide cannulae (Plastic One) were implanted in the ventrolateral OFC (anterioposterior +3.5mm, mediolateral ±2mm and dorsoventral +3.9mm; all coordinates are from bregma), secured with screws (Plastic One) and dental cement (Ortho-Jet, Lang Dental).

### Alcohol seeking and drug infusion

First, rats underwent IA-20%2BC paradigm for 7 weeks, as described above. Rats drinking more than 3.5 g/kg/24h on the last week of IA-20%2BC were selected for alcohol self-administration training in an operant self-administration (OSA) paradigm (37) with modification adapted from (21, 38, 39). Training was conducted in operant chambers (Med-Associates; Georgia, VT) using a fixed ratio 3 (FR3) (i.e. three active lever presses result in the delivery of one alcohol reward) during daily 30 minutes-session (Mon-Fri) as previously described (37). An inactive lever was also extended during the operant sessions but had no programmed consequence. Following stable baseline of alcohol responding, rats underwent stereotaxic surgery for cannula placement in the OFC, as described above. After a one-week recovery period, operant self-administration procedure was resumed and handling for drug microinfusions began. Rats received infusion of either vehicle or rapamycin (50ng/μl/side) 3 hours or R025-6981 (10μg/μl/side) 15 minutes before a single 30 minutes-extinction session. Drugs were injected with bilateral infusion needles (Plastics One) that projected 0.5 mm past the end of the guide cannula, at a rate of 0.5μl/minute.

Drug testing was performed using a “within-subject” design in which rats received both treatments in counterbalanced order.

Upon establishing a baseline of IA-20%2BC, a separate cohort of animals received an infusion of R025-6981 (10 μg/μl/side) or vehicle 15 minutes prior to the end of the last withdrawal session. The OFC was harvested at the end of the 24-hrs withdrawal session and were processed for western blot analysis.

### Habitual and goal-directed alcohol seeking and drug infusion

After 7 weeks of IA-20%2BC, high-drinking rats (alcohol intake >3.5 g/kg/24h) were trained to self-administer 20% alcohol in an operant self-administration (OSA) paradigm (37) with modification adapted from (21, 38, 39). Self-administration training was initiated under FR1 for 3 consecutive overnight sessions. Afterwards, the sessions lasted for an hour. Only one lever was available during the operant sessions. Rats were pseudo-randomly assigned to 2 groups and were subjected to a random interval (RI), or a random ratio (RR) schedule of reinforcement which biases responding towards habitual or goal directed actions, respectively (16). Only rats that self-administered alcohol more than 0.3g/kg/hour were included in the study.

RI training: During RI training, each reward was delivered following one lever press occurring after variable reward delivery time intervals (16). Three sessions were under RI-5 (i.e. inter-reward time was 5 seconds on average, ranging from 1 to 10 seconds), followed by three RI-15 sessions (i.e. inter-reward time was 15 seconds on average ranging from 10 to 30 seconds). Reinforcement was then shifted to RI-30 (i.e. inter-reward time was 30 seconds on average ranging from 10 to 50s) for at least 20 sessions (**Timeline**, Fig. 2a).

**Figure 1.**
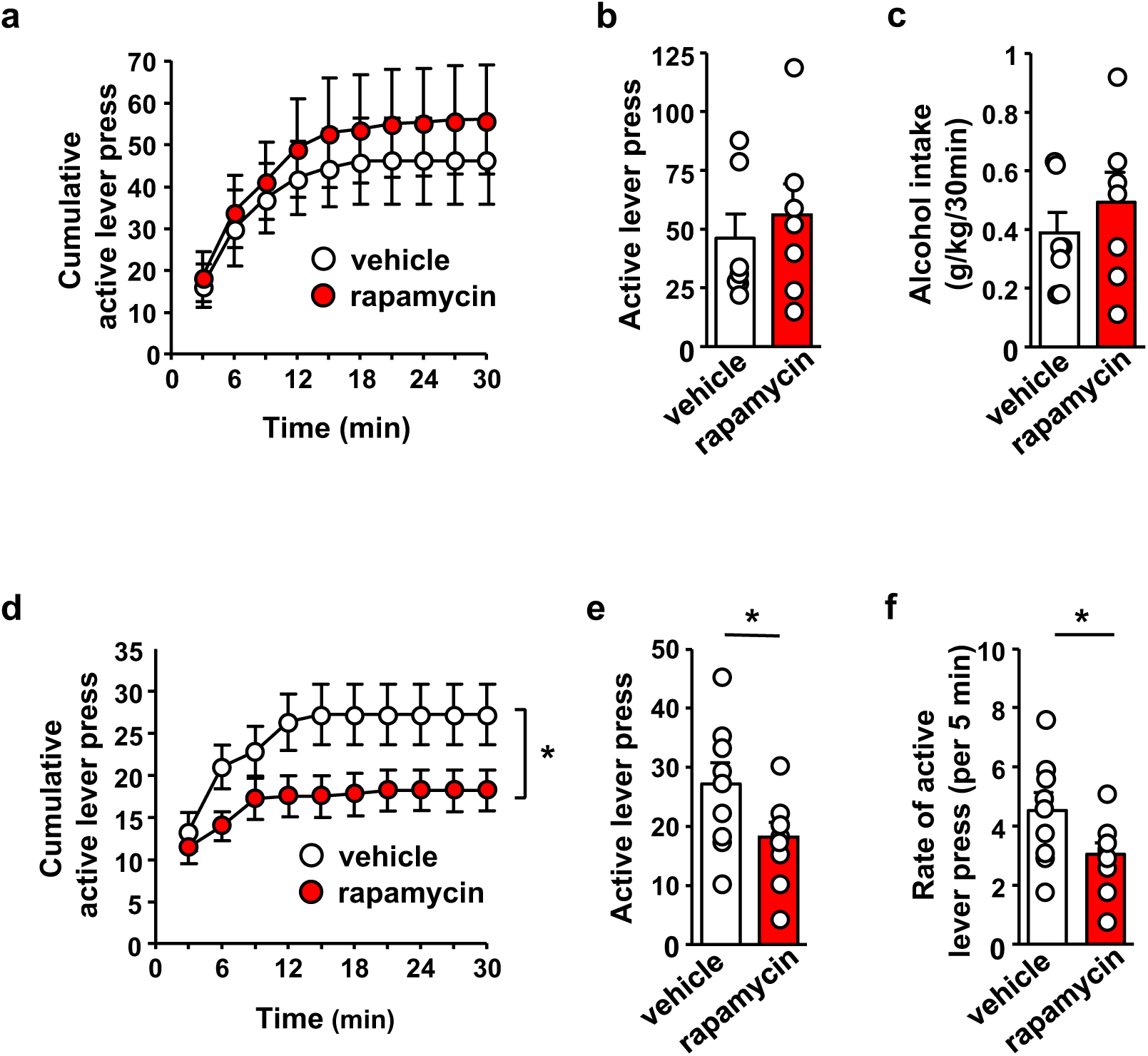
Inhibition of mTORC1 in the OFC reduces alcohol seeking. **(a-c) Intra-OFC infusion of rapamycin does not alter self-administration of alcohol.** Rats underwent 7 weeks of IA-20%2BC, and were then trained to self-administer 20% alcohol using a FR3 schedule. Vehicle (white) or rapamycin (50ng/μl, red) was infused bilaterally in the OFC 3 hours before a 30 min self-administration session. (**a**) RM ANOVA of cumulative lever presses did not identify a significant main treatment effect (F_1,12_=0.26, p>0.05). Two-tailed paired t-test revealed that the number of active lever presses (**b**) (t_6_=1.40, p>0.05), and amount of alcohol consumed (**c**) (t_6_=1.51, p>0.05) did not differ between treatment groups. **(d-f) Intra-OFC infusion of rapamycin inhibits lever presses during extinction.** Vehicle (white) or rapamycin (50ng/μl, red) was infused in the OFC 3 hours before a 30 min extinction session, and responses on the previously active lever were recorded. (**d**). RM ANOVA revealed a significant main treatment effect (F_1,16_=4.40, p=0.05) of cumulative lever presses. Two-tailed paired t-test revealed that the number (per 5 min) (**e**) t_8_=3.31, p<0.05, and rate of lever presses (**f**) (t_8_=3.30, p<0.05) were reduced in the rapamycin-treated animals. Data are presented as individual values and mean±SEM. *p<0.05. (a-c) n=7, (d-e) n=9.

**Figure 2.**
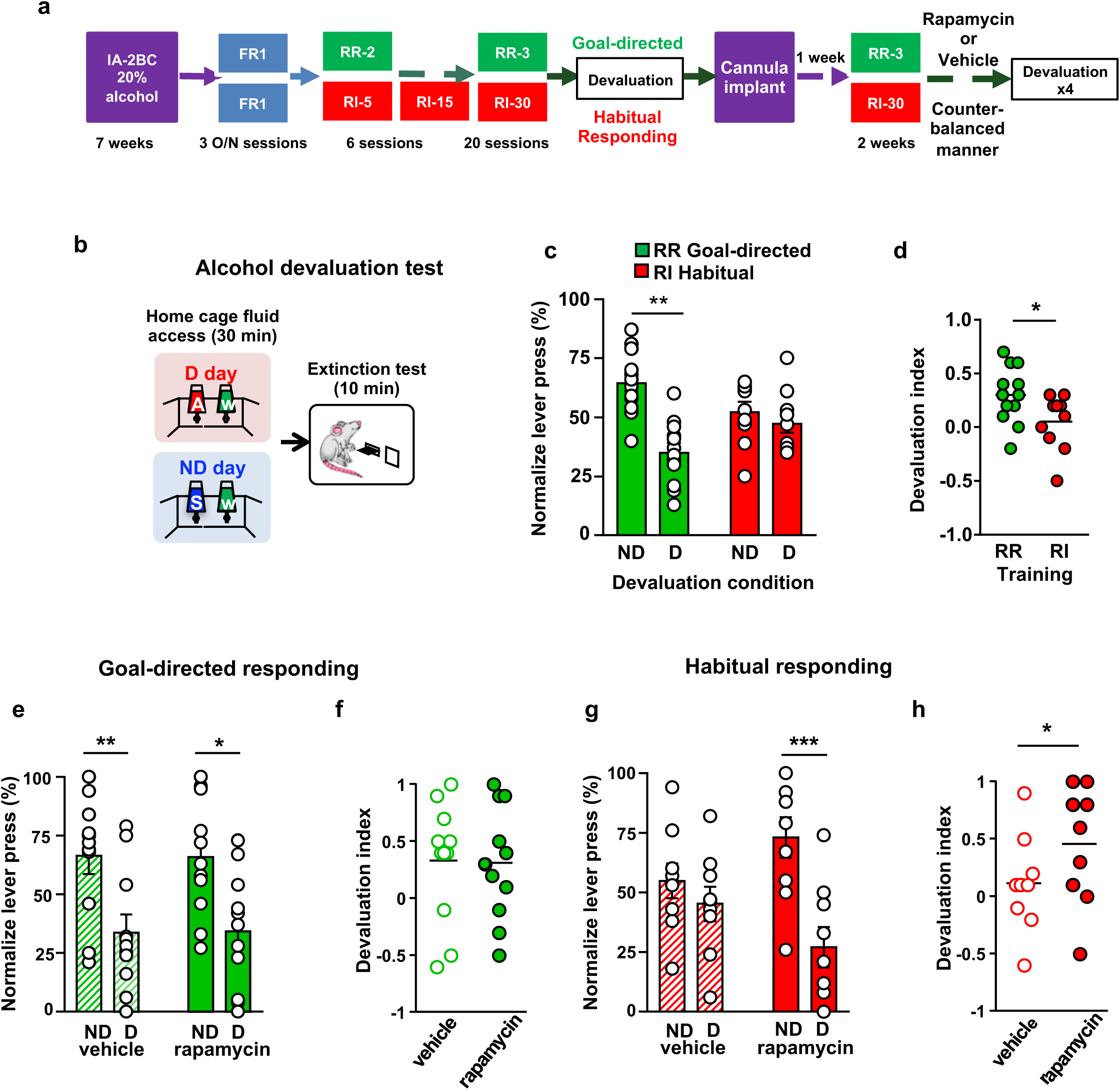
Inhibition of mTORC1 in the OFC attenuates habitual responding for alcohol. **(a) Timeline of experiment.** Rats underwent 7 weeks of IA20%-2BC. Rats were the trained to operant self-administer 20% alcohol during 3 overnight FR1 sessions and were pseudo-randomly assigned to 2 groups. One group was subjected to a progressive RI reinforcement schedule, while the other was assigned a RR schedule. Following a stable RI-30 or RR-3 responding, rats underwent alcohol devaluation testing depicted in **b**. Upon establishing the behavior, rats underwent stereotaxic surgery to bilaterally implant a guide cannula. One week later, RI-30 or RR-3 training was resumed for 2 weeks, after which rats received microinjections of vehicle or drug in a counter-balanced manner, prior to the devaluation test. **(b) Alcohol devaluation test.** On test day, rats were pre-fed with 20% alcohol for 30 minutes in their home cage during devalued day (D) or with 1% sucrose in the non-devalued day (ND), and normalized lever pressing (percent of total presses on D or ND, see methods) was used to evaluate goal-directed or habitual responding during a 10-minuted extinction session. **(c-d) RR trained animals are goal directed whereas RI trained animals show habitual behavior.** RR trained rats (green) press less during devalued day as compared to RI trained animals (red). Two-way ANOVA with Sidak post hoc analysis indicates a significant decrease in normalized lever pressing on D vs ND in RR-trained rats (p=0.0018), but not in RI-trained rats (p=0.8014) (Main effect of devaluation: F (1, 20) = 9.391, p=0.0061; reinforcement schedule X devaluation: F (1, 20) = 4.736, p=0.0417). **(c)** Devaluation index was calculated as the ratio of ND-D/ND+D. RR trained rats show a higher devaluation index as compared to the RI trained rats. Two-tailed unpaired t-test shows a significant difference in devaluation index (t_16_=3.266, p=0.0049). **(e-f) Intra-OFC infusion of rapamycin does not alter goal-directed alcohol seeking.** Rats that were trained using an RR schedule of reinforcement received an intra-OFC infusion of vehicle (hatched green) or rapamycin (50ng/μl) (green) 3 hours before the alcohol devaluation procedure as in (**b**), and the number of lever presses were measured during extinction. **(e)** Total lever press-based normalization indicates significant decreases in lever pressing between ND and D days in both vehicle-treated (p=0.0085) and rapamycin-treated (p=0.0117) RR-trained rats (Main effect of devaluation: F(1,40)=17.65, p=0.0001) **(f)** Devaluation index. No significant difference was detected (two-tailed paired t-test, t_13_=1.135, p=0.2768)**. (g-h) Intra-OFC infusion of rapamycin reduces habitual lever presses.** Rats that were trained using an RI schedule of reinforcement (red) received an intra-OFC infusion of vehicle (hatched red) or rapamycin (50ng/μl) (red) 3 hours before the alcohol devaluation procedure as in (**b**), and the number of lever presses were measured during extinction. **(g)** Total lever press-based normalization indicates significant decreases upon intra-OFC infusion of rapamycin- (p=0.0005), but not vehicle (p=0.6381) in RI trained rats on D compared to ND days (Main effect of devaluation: F(1,32)=12.38, p=0.0013; treatment X devaluation: F(1,32)=5.327, p=0.0276) **(h)** Devaluation index. Two-tailed paired t-test shows a significant difference in devaluation index (t_16_=3.557, p=0.0026). Data are presented as mean±SEM. *p<0.05, **p<.01 ***p<0.001. **(c-d)** n=8-10, **(e-f)** n=8-11, **(g-h)** n=8-9.

RR training: During RR training, reward was delivered following a variable number of lever responses (16). Six sessions under RR-2 (1 reward delivery following on average 2 lever presses with number of presses ranging from 1 to 3) followed by RR-3 (1 reward delivery following on average 3 lever presses with number of presses ranging from 1 to 5) for at least 20 sessions (**Timeline**, Fig. 2a).

Following stable RI-30 or RR-3 responding, rats underwent alcohol devaluation testing as described in (38) with modifications. Rats were given home-cage 2BC access to 1% sucrose on non-devalued (ND) days or 20% alcohol on devalued (D) days for 30 minutes (Fig. 2b). On each day, immediately after home-cage pre-feeding, rats underwent a 10-minutes extinction test during which unrewarded lever presses were recorded. A “within-subject” design was used in which ND or D sessions were in a counterbalanced order, with 2 standard retraining sessions between the tests. Rats that did not acquire the behavior and were pressing less than 5 lever presses per session on ND day were excluded from analysis. To minimize the effects of individual variance in total lever pressing we normalized ND and D lever presses (15, 16, 21). Normalized lever pressing was calculated as ND or D lever presses/total number of lever presses, respectively. Normalizing devaluation lever presses produces a distribution of data, where 50% is equivalent to equal lever pressing between ND and D (16). Devaluation index was calculated as the ratio of lever presses on ND day–lever presses on D day/total number of lever presses. At the end of the training period and upon acquiring the behavior (RR, goal directed behavior and RI, habitual behavior), rats underwent stereotaxic surgery to bilaterally implant a guide cannula in the ventrolateral OFC, as described above. One week later, RI-30 or RR-3 training was resumed for 2 weeks after which rats received microinjections of either vehicle or rapamycin (50ng/μl/side) 3 hours or R025-6981 (5μg/μl/side) 15 minutes prior an alcohol devaluation testing.

### Habitual sucrose seeking

RI30 training was similar to the procedure described for alcohol except that 8% sucrose was initially used and progressively reduced to 1% across training sessions. Only ten RI30 sessions (2 weeks) were conducted prior to the initial outcome devaluation test. Rats were then implanted with a guide cannula in the OFC as described above. After one-week recovery period, sucrose self-administration was resumed for two weeks, after which rats received microinjections of vehicle or drug in a counter-balanced manner, prior to the devaluation test. Sensitivity to changes in sucrose value was tested using a sucrose devaluation procedure which was similar to alcohol devaluation except that rats had 2BC access to 20% alcohol on ND day and 1% sucrose on D day (**Timeline**, Fig. 3a-b).

**Figure 3.**
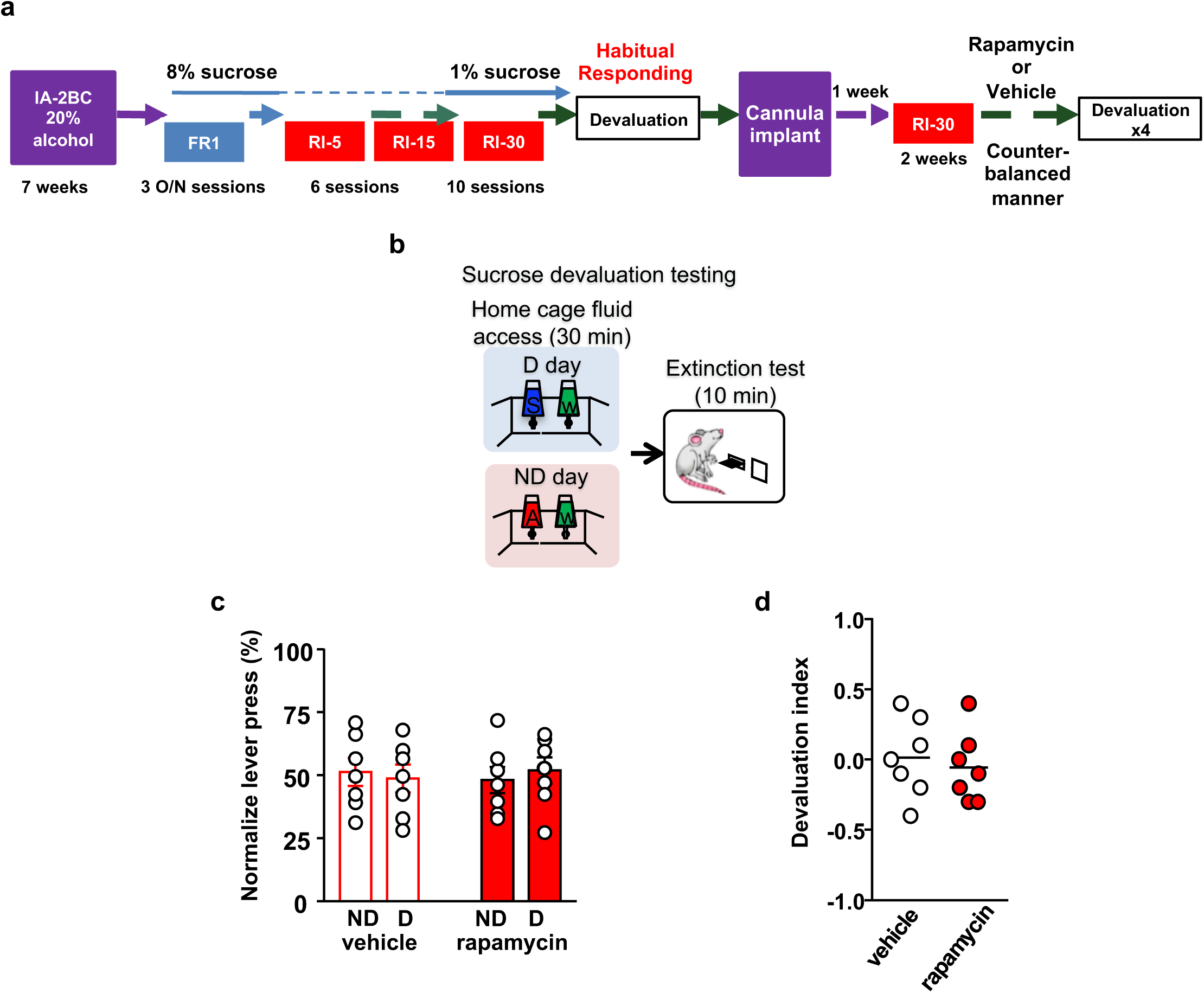
Habitual sucrose responding is not affected by mTORC1 inhibition in the OFC. **(a) Timeline of experiment.** Rats underwent 7 weeks of IA20%-2BC, and were then trained to operant self-administer 8% sucrose. Sucrose concentration was progressively reduced to 1% during 3 overnight FR1 sessions. Rats were then subjected to an RI reinforcement schedule. Following a stable RI-30 responding, rats underwent alcohol devaluation testing depicted in **b**. Upon establishing the behavior, rats underwent stereotaxic surgery to bilaterally implant a guide cannula. One week later, RI-30 training was resumed for 2 weeks, after which rats received microinjections of vehicle or rapamycin (50ng/μl) in a counter-balanced manner, prior to the devaluation test. **(b) Sucrose devaluation test.** Habitual behavior was probed in an identical way to the alcohol-trained cohorts with the exception that sucrose pre-feeding was considered the devaluing context and alcohol was the non-devalued substance. **(c)** Two-way RM ANOVA failed to show a significant effect of repeated testing (Main effect of devaluation: F(1,12)=0.006464, p>0.9999; treatment X devaluation: F(1,12)=0.7257, p=0.4110) on lever presses between devalued and non-devalued days when habitually responding animals were pretreated with either vehicle or rapamycin. n=7 **(d)** Devaluation index. Two-tailed paired t-test detected no significant difference between the treatment groups (t=0.7024, p=0.5087). n=7.

### Histology

For verification of cannula placement, brains were post-fixed in 4% PFA for one week, then rapidly frozen and sectioned into 50μm coronal slices using a Leica CM3050 cryostat (Leica Biosystems). Slices were stained using cresyl violet and examined for cannula placement using a bright-field microscope. All rats had correct guide cannula placement and were included in the study.

### *Ex vivo* NMDA treatment

Naïve rats were euthanized by isoflurane and decapitated. Brains were removed and sectioned into 300 µm-thick slices at 4° C in a solution of aCSF containing (in mM) 200 sucrose, 1.9 KCl, 1.4 NaH_2_PO^4^, 0.5 CaCl_2_, 6 MgCl_2_, 10 glucose, 25 NaHCO_3_, 4 ascorbic acid; 310-330 mOsm. Immediately after sectioning, OFC-containing slices were recovered in aCSF (in mM: 125 NaCl, 2.5 KCl, 1.4 NaH_2_PO_4_, 25 NaHCO_3_, 2 CaCl_2_, 1.3 MgCl_2_, 25 glucose, 0.4 ascorbic acid) heated at 32° C for 30 minutes and then moved to an aCSF solution containing 1 μM TTX at room temperature for 30 minutes (40). The OFC was dissected and incubated for 30 minutes in aCSF containing 1 μM TTX at room temperature. OFC sections were then treated for 3 minutes with vehicle (0.1% DMSO) or NMDA (25μM) in TTX-containing aCSF, and western blot analysis was conducted as described below.

### Western blot analysis

Tissue were homogenized in ice-cold radio immunoprecipitation assay (RIPA) buffer (in mM: 50 Tris-Cl, 5 EDTA, 120 NaCl, 1% NP-40, 0.1% deoxycholate, 0.5% SDS, protease and phosphatase inhibitors), using a sonic dismembrator. Protein content was determined using BCA kit. Tissue homogenates (30 μg per sample) were resolved on NuPAGE 10% Bis-Tris gels at 100 V for 2 hours and transferred onto nitrocellulose membranes at 30V for 2 hours. Blots were blocked with 5% milk-phosphate-buffered saline with 0.1% tween-20 at room temperature and then probed with primary antibodies overnight at 4°C. Membranes were washed and probed with HRP-conjugated secondary antibodies for 2 hours at room temperature, and bands were visualized using ECL. Band intensities were quantified by ImageJ (National Institutes of Health, MD, USA).

### Data analysis

All data are expressed as the mean ± SEM. GraphPad Prism 7.0 (GraphPad Software, Inc., La Jolla, CA, USA) was used to plot and analyze the data. D’Agostino–Pearson normality test and *F*-test/Levene test were used to verify the normal distribution of variables and the homogeneity of variance, respectively. The sample sizes were based on previous publications (8, 41). Data between two groups were compared using two-tailed paired or unpaired *t*-test. Data from multiple groups against one group were compared using a two-way repeated-measures analysis of variance followed when appropriate by a Tukey multiple comparisons test or a Sidak’s multiple comparisons test. Statistical significance was set at p<0.05.

## RESULTS

### mTORC1 in the OFC contributes to alcohol seeking

We first examined whether activation of mTORC1 in the OFC contributes to the maintenance of heavy alcohol use. Rats that underwent 7 weeks of IA-20%2BC received an intra-OFC bilateral infusion of vehicle or the mTORC1 inhibitor, rapamycin (50ng/μl), 3 hours prior to a drinking session (8) (Fig. 1 - Supplement 1). As shown in Fig. 1 - Supplement 2, intra-OFC administration of rapamycin did not impact voluntary intake of alcohol. Next, rats that underwent IA-20%2BC for 7 weeks were trained to self-administer 20% alcohol in an operant self-administration (OSA) paradigm (42). Upon establishing a baseline of alcohol lever presses, rats received a bilateral infusion of vehicle or rapamycin (50ng/μl) into the OFC 3 hours prior to an OSA session, and the number of lever presses and alcohol consumption were measured (8). As shown in Fig. 1a-c, inhibition of mTORC1 in the OFC failed to alter the number of lever presses (Fig.1a-b), and alcohol consumed (Fig. 1c). Inactive lever pressing (Fig. 1 - Supplement 3a). The latency to the first or last lever press were also unaltered (Fig. 1 - Supplement 3b-c). Next, we determined whether mTORC1 plays a role in alcohol seeking by testing whether rapamycin alters lever presses during an extinction session. Intra-OFC administration of rapamycin suppressed both the number (Fig. 1d-e), and the rate (Fig. 1f) of rats’ responding on the lever that was previously associated with alcohol. The reduction in alcohol seeking was not due to alterations in locomotion, as inter-response intervals were similar in vehicle and rapamycin-treated rats (Fig. 1 - Supplement 4). Together, these data suggest that mTORC1 in the OFC does not contribute to alcohol drinking *per se,* but does promote alcohol seeking.

### mTORC1 in the OFC participates in habitual responding for alcohol

Reward seeking can be driven by goal-directed or habitual compulsive behaviors (43–45). We next examined whether mTORC1 in the OFC plays a role in goal-directed or habitual alcohol seeking using a satiety outcome devaluation procedure. Rats that first underwent 7 weeks of IA-20%2BC were then trained to lever press for 20% alcohol using either an random ratio (RR) or random interval (RI) schedule of reinforcement, which biases lever responding toward goal-directed (RR) or habitual (RI) actions (16, 21, 38) (**Timeline**. Fig. 2a). RR and RI trained rats showed a similar number of presses throughout the training (Fig. 2 - Supplement 1a), however alcohol consumption was slightly higher during RI training (Fig. 2 - Supplement 1b), presumably due to fewer lever presses being required to obtain the same volume of alcohol, compared with RR training. Rats trained to self-administer alcohol in a goal-directed manner are sensitive to outcome devaluation, while habitually-trained rats are not, e.g. animals who habitually press a lever to obtain a reward will continue to press that lever, even when the relative value of the reward is reduced due to satiety (16, 21, 38, 46). Thus, we conducted a devaluation procedure to differentiate between goal-directed and habitual responding, (Fig. 2b). On the devalued (D) day, rats had 30 minutes of home-cage access to 20% alcohol, the reward previously associated with lever responding, and on the non-devalued (ND) day, rats had 30 minutes of home-cage access to 1% sucrose, a reward not previously associated with lever responding (Fig. 2b). Rats then underwent a 10-minute extinction test, during which lever presses were recorded (Fig. 2b). In order to minimize the effects of individual variability in overall lever pressing, and to graphically represent the distribution of lever presses between ND and D, we normalized lever-pressing data such that 50% indicates identical lever pressing on both days as shown in (15, 16, 21). As shown in Fig. 2c-d, rats trained on an RR schedule exhibit a decrease in responding following alcohol devaluation, while RI-trained rats remained insensitive to devaluation. These data show that RI training can produce habitual responding for alcohol in rats.

Next, to test whether mTORC1 in the OFC participates in goal-directed or habitual alcohol seeking, we infused rapamycin (50ng/μl) or vehicle bilaterally, 3 hours prior to a devaluation session, and examined lever presses in RR- and RI-trained groups (Fig. 2a, Fig. 2 - Supplement 2). Rapamycin administration did not alter home cage alcohol intake prior to the extinction session in either group (Fig. 2 - Supplement 3). Intra-OFC administration of rapamycin did not impact responding in RR-trained (goal-directed) cohorts, as lever presses were reduced in rats that were pre-fed with alcohol and treated with either vehicle or rapamycin (Fig. 2e-f). In contrast, inhibition of mTORC1 in the OFC reduced responding in the RI-trained (habitual) rats in the devalued condition (Fig.2g-h). Critically, the reduction in responding in the rapamycin-treated RI-trained rats paralleled the responding of RR-trained animals (Fig.2e-f vs. g-h). These data suggest that mTORC1 inhibition in the OFC restores goal-directed alcohol seeking in animals trained to habitually self-administer alcohol.

To determine whether mTORC1 in the OFC plays a role in habitual responding to a natural reward, animals were trained to self-administer sucrose under an RI schedule of reinforcement, and lever presses were measured in rats treated with rapamycin (50ng/μl) or vehicle (Fig. 3 - Supplement 1) during an extinction session in animals that were first offered sucrose (devalued) or alcohol (non-devalued) in their home cage (**Timeline**, Fig. 3a-b). Rapamycin administration did not alter home cage sucrose intake prior to the extinction session (Fig. 3 - Supplement 2) As shown in Fig. 3c-d, habitual responding for sucrose was similar in the vehicle- and rapamycin-treated animals. Together, these data suggest that mTORC1’s contribution to habitual behavior is not generalized to all rewarding substances.

### Alcohol-dependent mTORC1 activation in the OFC requires GluN2B

Next, we set out to elucidate the mechanism by which long-term heavy alcohol use activates mTORC1 in the OFC. Glutamatergic inputs from cortical and subcortical structures project to the OFC (47, 48). Extracellular glutamate content is elevated in cortical regions of humans and rats during alcohol withdrawal (49). N-methyl D-Aspartate receptors (NMDAR) are targets of alcohol (50). For example, GluN2B-containing NMDARs are activated by alcohol in the hippocampus and dorsal striatum (50). Stimulation of GluN2B activates the guanine nucleotide releasing factor 1 (GRF1), the activator of H-Ras (51) (Figure 4 – Supplement 1). Since H-Ras is upstream of mTORC1 (52) (Figure 4 – Supplement 1), we hypothesized that alcohol-dependent stimulation of GluN2B activates mTORC1 in the OFC. We first tested whether NMDAR stimulation activates mTORC1. To do so, OFC slices were briefly treated with NMDA (25μM), and the activation of CaMKII and mTORC1 were tested by measuring the phosphorylation of mTORC1 downstream targets, S6K and S6 (Figure 4 – Supplement 1). We found that *ex vivo* treatment of the OFC with NMDA led to increased phosphorylation, and thus activation, of CaMKII, a marker of NMDAR activation (Fig. 4a-b), as well as of mTORC1 (Fig. 4a-b). These data suggest that NMDAR stimulation activates mTORC1 in the OFC.

**Figure 4.**
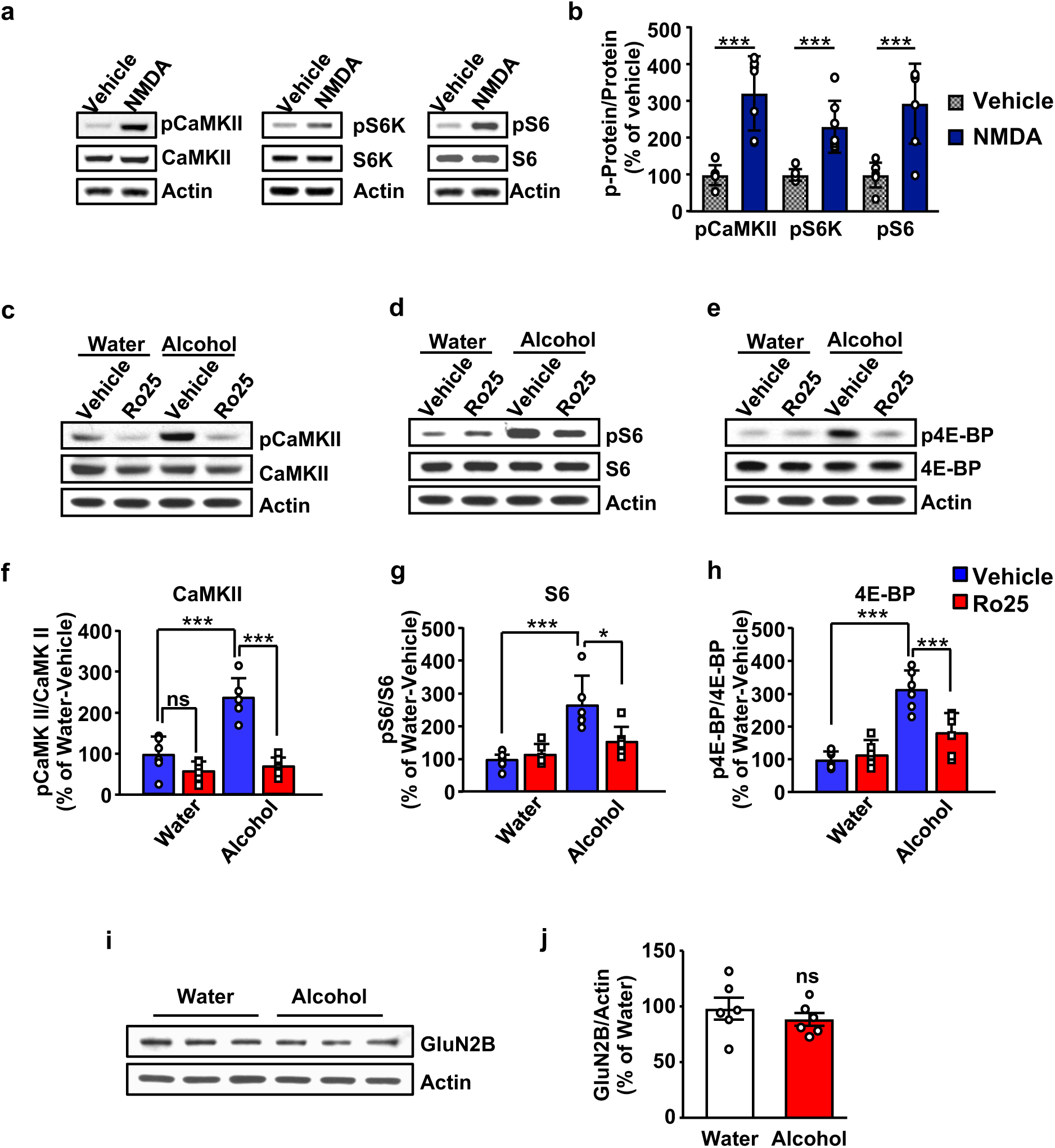
GluN2B is required for mTORC1 activation by alcohol. **(a-b) Stimulation of NMDA receptors activates mTORC1 in the OFC.** Vehicle (0.1% DMSO) or NMDA (25μM) was applied to OFC slices of naïve rats for 3 minutes. Phosphorylation of CaMKII, S6K and S6 were determined by western blot analysis. Total levels of the proteins were measured as well, and actin was used as a loading control. **(a)** Representative blots of CaMKII (left), S6K (middle), and S6 (right) phosphorylation in OFC slices treated with vehicle or NMDA. **(b)** ImageJ was used for optical density quantification. Data are expressed as the average ratio +/-S.E.M of phospho-CaMKII to CaMKII, phospho-S6K to S6K and phospho-S6 to S6, and are expressed as percentage of vehicle. Significance was determined using two-tailed unpaired *t*-test. (CaMKII: t_12_=5.61, p<0.001; S6K: t_12_=4.77, p<0.001; S6: t_12_=4.51, p<0.001). **(c-h) Alcohol-dependent mTORC1 activation is attenuated by the GluN2B inhibitor, Ro25-6981.** Rats underwent 7 weeks of IA-20%2BC resulting in an average of alcohol intake of 4.37 g/kg/24 hours. Rats received an intra-OFC bilateral infusion of vehicle (saline) or Ro25-6981 (Ro25, 10μg/μl), 15 minutes prior to the beginning of the water only session and the OFC was dissected 24 hours later. Water only consuming rats were used as controls. (**c**-**e**) Representative images of CaMKII (**c**), S6 (**d**) and 4E-BP (**e**) phosphorylation in water vs. alcohol-exposed animals that were pre-treated with vehicle or Ro25-6981. (**f**-**h**) ImageJ was used for optical density quantification. Data are expressed as the average ratio +/- S.E.M of phospho-CaMKII to CaMKII (**f**), phospho-S6 to S6 (**g**), and phospho-4E-BP to 4E-BP (**h**), and are expressed as percentage of water + vehicle group. Two-way RM ANOVA showed a significant effect of alcohol on pCaMKII (F_1,20_=24.66, p<0.0001), pS6 (F_1,20_=21.92, p=0.0001), and p4E-BP (F_1,20_=48.23, p<0.0001). In addition, two-way RM ANOVAs indicated a main effect of drug treatment for pCaMKII (F_1,20_=48.51, p<0.0001), pS6 (F_1,20_=4.58, p<0.05), and p4E-BP (F_1,20_=8.49, p<0.01), as well as treatment x alcohol interaction for pCaMKII (F_1,20_=18.57, p<0.01), pS6 (F_1,20_=7.559, p<0.05), and p4E-BP (F_1,20_=13.22, p<0.01). Tukey’s multiple comparison post hoc analysis revealed that alcohol increased pCaMKII (p<0.001), pS6 (p<0.001), and p4E-BP (p<0.001) in vehicle-treated animals and that pretreatment with R025-6981 reduced the phosphorylation of pCaMKII (p<0.0001), S6 (p<0.05), and 4E-BP (p<0.001). **(i-j**) **Chronic alcohol intake does not alter the level of GluN2B in the OFC.** At the conclusion of the 7 weeks of IA-20%2BC, the OFC was collected from rats at the end of the last 24 hours water session. GluN2B levels were determined by western blot analysis, and actin was used as a loading control. **(i)** Representative blots of GluN2B protein levels in the OFC of alcohol consuming vs. water only consuming rats. **(j)** ImageJ was used for optical density quantification. Data are expressed as the average ratio +/-S.E.M of GluN2B levels, expressed as percentage of water. Significance was determined using two-tailed unpaired t-test. Alcohol had no effect on GluN2B levels (t_10_=0.85, p>0.05). *p<0.05, ***p<0.001. (**a-b)** n = 7 per group, (**c-j**) n = 6 per group.

We previously showed that mTORC1 is activated in the OFC in response to a binge drinking session which persists for at least 24 hours into alcohol abstinence (1). Thus, we next tested whether GluN2B is required for alcohol-dependent mTORC1 activation. To do so, rats that underwent 7-weeks of IA-20%2BC paradigm or that drank water only for the same duration of time, received a bilateral intra-OFC infusion of the selective GluN2B antagonist Ro-25-6981 (53) (10μg/ul) or vehicle, 15 minutes prior to the end of the last alcohol drinking session, and mTORC1 activation was examined at the end of the last 24 hour water only session by measuring the phosphorylation of mTORC1 downstream targets, 4-EBP and S6 (Fig. 4 - Supplement 1, **Timeleine** Figure 4 - Supplement 2). In vehicle-treated rats, alcohol produced a robust activation of CaMKII (Fig. 4c,f) and mTORC1, as measured by S6 (Fig. 4d,g), and 4-EBP phosphorylation (Fig. 4e,h). In contrast, R025-6981-treated rats exhibited a marked reduction in both CaMKII and mTORC1 activation during alcohol withdrawal Fig. 4c-h. In contrast, GluN2B protein levels in the OFC were unchanged by alcohol (Fig. 4i-j). Together our data suggest that GluN2B mediates mTORC1 activation following alcohol consumption.

### GluN2B in the OFC is required for alcohol seeking and habit

We reasoned that, if GluN2B is upstream of mTORC1, then GluN2B should also participate in habitual alcohol seeking. First, we examined whether inhibition of GluN2B alters alcohol seeking during extinction. We found that intra-OFC administration of Ro25-6981 (10μg/μl) 15 minutes prior to an extinction session (Fig. 5 - Supplement 1) suppressed alcohol seeking, as indicated by a reduction in cumulative (Fig. 5a), active lever presses (Fig. 5b), as well as the rate of lever pressing (Fig. 5c). The reduction of alcohol seeking by Ro25-6981 was not due to alteration of locomotion as inter-response intervals were similar in vehicle-treated and Ro25-6981-treated animals (Fig. 5 - Supplement 2).

**Figure 5.**
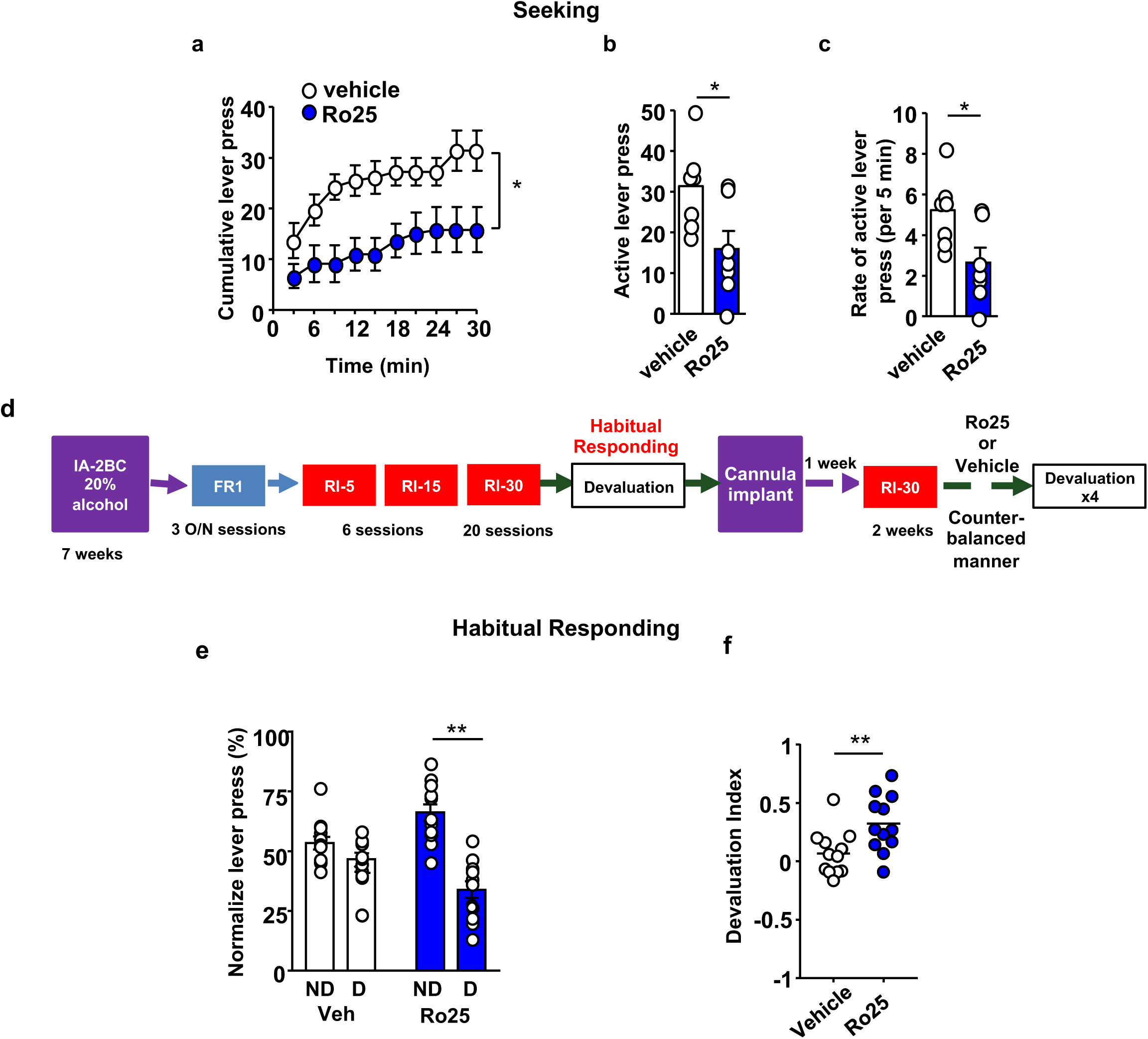
GluN2B in the OFC promotes alcohol seeking and habitual alcohol responding. (**a-c**) **Intra-OFC administration of Ro25-6981 decreases alcohol seeking.** Rats underwent 7 weeks of IA-20%2BC, and were then trained to self-administer 20% alcohol. Vehicle (white) or Ro25-6981 (Ro25, 10μg/μl, blue) was infused bilaterally in the OFC 15 minutes prior to a 30 minutes extinction session. **(a)** Cumulative lever presses. Two-way RM ANOVA revealed a significant main treatment effect (F_1,12_=10.89, p<0.05). Two-tailed paired t-test revealed that the number **(b)** and rate (per 5 minutes) **(c)** of lever press were significantly different in the vs. vs. Ro25-6981 groups ng (both t_6_=2.62, p<0.05). **(d) Timeline of experiment.** Rats underwent 7 weeks of IA20%-2BC, and were then trained to operant self-administer 20% alcohol using a progressive RI reinforcement schedule. Following a stable RI-30 lever presses, rats underwent stereotaxic surgery to bilaterally implant guide cannulae in the OFC. One week later, RI-30 training was resumed for 2 weeks, after which rats received microinjections of vehicle (white) or Ro25-6981 (5 μg/μl, blue) in a counter-balanced manner, 15 minutes prior to a 30 minutes home cage 20%2BC alcohol drinking session, and the number of lever presses were measured during extinction. **(e-f) Intra-OFC administration of Ro25-6981 attenuates habitual alcohol seeking**. (**e**) Two way RM ANOVA showed a main effect of devaluation (F_1,33_=13.72, p<0.001) as well as a treatment x devaluation interaction (F_1,33_=6.31, p<0.05). Sidak’s multiple comparison test detected a significant difference for Ro25-6981 (p<0.001) on D compared to ND days. **(f)** Devaluation index. Two-tailed paired t-test indicated a significant difference (t_15_=3.72, p<0.01). Individual results and mean±SEM are shown, (**a-c**) n=7, (**e-f**) n=12. *p<0.05, ** p<0.01.

Finally, we tested whether GluN2B in the OFC contributes to habitual alcohol seeking. The timeline of the experiment is depicted in Fig. 5d. Intra-OFC infusion of R025-6981 (5μg/μl) did not impact home cage alcohol consumption in a 2BC test during devalued days (Fig. 5 - Supplement 3). In contrast, administration of Ro25-6981 (5μg/μl) into the OFC reverted habitual responding for alcohol to goal directed responding (Fig. 5e-f). Together, these data indicate that the recruitment of GluN2B signaling by alcohol is required for alcohol seeking and habit.

## DISCUSSION

Using operant training protocols that bias alcohol seeking towards habitual or goal-directed behavior, we present data implicating mTORC1 in the OFC in mechanisms underlying the development and/or maintenance of habitual behavior associated with alcohol use. In addition, we show that GluN2B in the OFC is required for alcohol-dependent mTORC1 activation and function. Together, our data suggest that the GluN2B/mTORC1 axis acts as a molecular switch that converts the OFC from driving goal directed to habitual alcohol use.

We found that inhibition of mTORC1 in the OFC attenuates lever pressing during extinction, suggesting that mTORC1 regulates alcohol seeking. The OFC has been implicated in updating reward anticipation based on the value of an expected outcome (17, 54), and O’Doherty et al. showed that the lateral OFC (lOFC) is activated during reward anticipation in humans (55). Thus, it is plausible that one of the roles of mTORC1 in the OFC is the strengthening of alcohol reward anticipation, thus promoting alcohol seeking.

We further show that blockade of mTORC1 activity in the OFC converts habitual alcohol responding to a more goal directed behavior. Habitual behavior is thought to rely on the strengthening of the dorsolateral striatum (DLS) over dorsomedial striatum (DMS), a brain region which participates in goal-directed behaviors (38, 56, 57). Renteria and colleagues showed that alcohol vapor exposure generates habitual behavior via the disruption of OFC to DMS circuit (39). Thus, it is plausible that mTORC1 activation is strengthening OFC projections to the DLS, and/or is weakening OFC to DMS projections. One caveat to consider, however, is that we measured habitual vs. goal directed responding for alcohol, whereas Renteria et al. measured responding for food. It is plausible that Renteria et al. tested whether alcohol alters decision making in general, while the current study is focused on habitual responding specifically for alcohol. It would be of interest to determine the normal role of mTORC1 in the OFC, and whether mTORC1 suppresses other decision-making tasks.

We previously showed that mTORC1 activation is detected in the lOFC (1), which was targeted in the current study. However, the structure of the OFC is complex. The medial OFC (mOFC) and lOFC are each part of anatomically and functionally distinct corticostriatal circuits in humans (58) and rodents (59). The mOFC has been reported to play a larger role in determining, updating, and storing the relative reward values of stimuli and actions (60). The lOFC, on the other hand, is proposed to be more involved in encoding the relationship between stimuli/actions and outcomes (60), reward anticipation and goal directed behavior (16). As mTORC1’s activation is localized to the lOFC, more research is required to determine if mTORC1 contributes to other lOFC-related behaviors.

The OFC has been associated with compulsive behavior (22), and recently Pascoli et al. reported that OFC neurons projecting to the dorsal striatum are associated with compulsive lever pressing in response to optogenetic activation of dopaminergic ventral tegmental area neurons (23). Compulsive drug use is associated with maladaptive habitual responding (57, 61), and we and others, provided animal data to support the notion that alcohol consumption can become compulsive (44, 62, 63). Thus, it is possible that mTORC1 signaling in the OFC is driving both habitual and compulsive alcohol seeking.

Interestingly, mTORC1 inhibition in the OFC does not alter habitual responding for sucrose, indicating that mTORC1 plays a specific role in the formation of habits in response to alcohol exposure and not natural rewards. The difference between the consequences of mTORC1 inhibition on habitual alcohol vs. sucrose seeking is striking, but it is important to note that, unlike alcohol, voluntary sucrose intake does not activate mTORC1 in the OFC (1).

Corbit et al. showed that RR-trained rats exhibit habitual alcohol self-administration after 8 weeks of training on an RR3 schedule of reinforcement (38). In the current study, rats undergoing RR training demonstrated persistent goal-directed alcohol seeking behaviors. The difference between the two studies is most likely to be due to the length of RR training (long in Corbit et al. (38) vs. short herein). It is possible that further training would drive our RR-trained goal-directed rats to exhibit more habitual phenotypes. Further research would be required to compare the models more accurately.

We previously reported that repeated cycles of binge alcohol drinking and withdrawal activates mTORC1 in the OFC (1), and we demonstrate herein that mTORC1 activation in the OFC depends, at least in part, on GluN2B suggesting that GluN2B is activated in response to alcohol use. The NMDAR is a target of alcohol (50), and alcohol enhances GluN2B phosphorylation in brain regions such as the cerebellum, the DMS and the hippocampus (50). GluN2B phosphorylation enhances the activity of the channel (69). Thus, it is plausible that alcohol-dependent GluN2B phosphorylation in the OFC enhances GluN2B function thereby enabling mTORC1 activation. Alternatively, the hyperactivity of GluN2B could be a compensatory homeostatic plasticity resulting from the inhibition of NMDAR by alcohol (50). In contrast to our findings that alcohol does not alter the expression of GluN2B in the OFC, Nimitvilai and colleagues showed that chronic exposure to alcohol vapor decreases in the expression of GluN2B (28). One important difference between our study and Nimitvilai et al. as well as additional reports on the interaction between the OFC and alcohol discussed herein, relates to the use of contingent vs. non-contingent alcohol administration. Specifically, prior to the OSA, rats were subjected to 7 weeks of IA-20%2BC. The IA-20%2BC paradigm models “problem drinkers” e.g. human subjects that suffer from AUD phenotypes such as binge drinking, compulsive drinking and craving (64). In our case, rats achieve blood alcohol concentrations (BAC) of 80 mg/dl (65) a BAC equivalent to binge drinking humans (66). In contrast, the majority of other reports described herein involved passive exposure of alcohol vapor, which achieves BAC of 150-250 mg/dl, and leads to physical dependence on alcohol (67). Humans who are dependent on alcohol show profoundly degraded white matter and reduced neuronal density in the OFC (31, 32), possibly an indication of alcohol-induced damage to this structure. Thus, it is likely that cycles of binge drinking do not damage the OFC which may be the case in animal models of alcohol dependence. As only a small percentage of alcohol users exhibit physical dependence (68), we believe that our findings have important implications for the understanding of the mechanisms underlying problem drinking and AUD.

What could be the mechanism by which mTORC1 contributes to alcohol seeking and habit? Activation of mTORC1 triggers the translation of a subset of mRNAs to protein in the cell body and in dendrites (3, 70). We previously showed that long-term excessive alcohol intake and reinstatement of alcohol place preference initiate the translation of several synaptic proteins in the NAc (11, 12, 36). We also found that reconsolidation of alcohol reward memories increases the immunoreactivity of mTORC1 targets in the OFC (41). Thus, it is possible that the consequences of mTORC1 activation in the OFC is the translation of synaptic proteins.

We show here that GluN2B activation in the OFC is sufficient to influence the formation and/or maintenance of alcohol seeking and habit. In line with the possibility that GluN2B activation contributes to the development of habit, DePoy et al. showed that habitual behavior, developed upon subchronic cocaine treatment during adolescence (71), is inhibited by the treatment of animals with the GluN2B inhibitor ifenprodil (71). DePoy et al. further showed that cocaine exposure during adolescence produces structural changes in the OFC which were attenuated by ifenprodil (71). While our data suggest that GluN2B in the OFC participates in behavioral inflexibility, Brigman et al. reported that GluN2B in the OFC plays a role in choice learning (72), It is plausible that alcohol-dependent neuroadaptations tilt the function of GluN2B from choice learning, e.g. a goal directed behavior, to habitual responding.

Gremel et al. showed that the endocannabinoid/CB1 receptor system in in OFC to DLS circuit promotes habit formation by weakening the OFC to DMS circuitry required for goal directed behavior (21). Interestingly, cannabinoid signaling has been shown to activate mTORC1 in hippocampal neurons (73). Furthermore, THC-dependent impairment of novel object recognition was reversed by the mTORC1 inhibitor, rapamycin (73). Thus, it is possible that endocannabinoids and mTORC1 are part of the same signaling cascade that gates goal-directed behaviors, thereby promoting habit. Finally, Gourley and colleagues reported that the brain-derived neurotrophic factor (BDNF) in the OFC participates in goal-directed behaviors (18, 74, 75). BDNF and mTORC1 play opposing roles in AUD (76). For example, BDNF keeps alcohol intake in moderation, and mTORC1 is associated with excessive alcohol use (76). As breakdown of the BDNF signaling cascade results in compulsive alcohol use (62). It is tempting to speculate that the balance between BDNF and mTORC1 signaling in the OFC determines whether alcohol use is kept in moderation or becomes habitual.

## Acknowledgements

ACKNOWLEDGEMENTS AND DISCLOSURES

This research was supported by the National Institute of Alcohol Abuse and Alcoholism, P50 AA017072 (D.R.) and R01 AA027474 (D.R.).

The authors do not have a financial or non-financial competing interests.

## FIGURE LEGENDS

**Figure 1 – Supplement 1.**
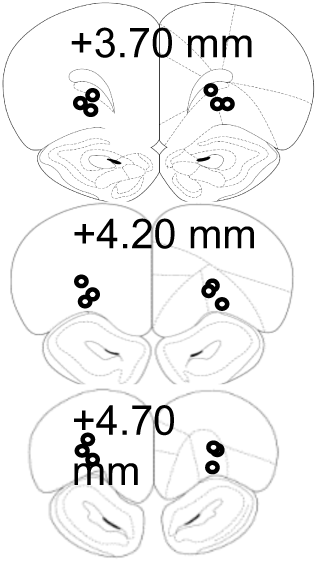
Schematic drawing of cannulae placement. Schematic illustration of coronal sections of the rat brain showing approximate bilateral placements of cannulae in the OFC according to Paxinos and Watson 2007.

**Figure 1 – Supplement 2.**
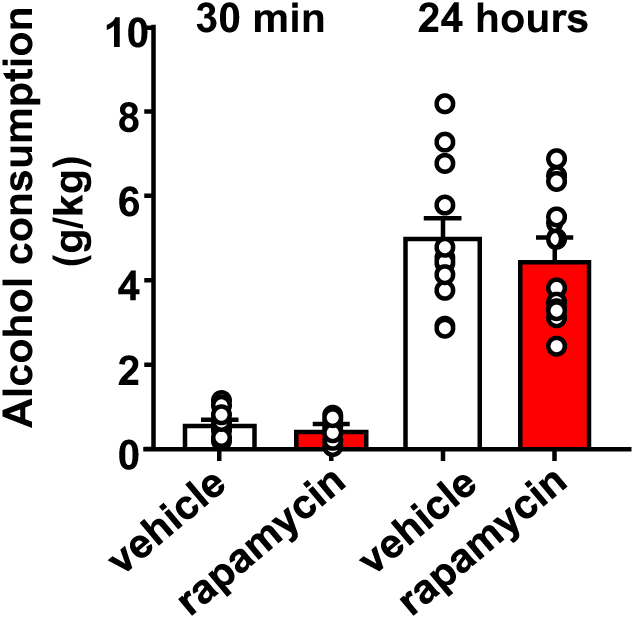
Inhibition of mTORC1 in the OFC does not alter alcohol consumption in a 2-bottle choice paradigm. Rats with a history of IA-20%2BC were infused with vehicle (white) or rapamycin (50ng/μl, red) into the OFC 3 hours before the beginning of the final 2BC session, and alcohol consumption was recorded at the 30 minutes and 24-hour time points. Significance between treatment groups was determined using RM ANOVA. No differences were observed in alcohol consumption at the 30 minutes (t_11_=1.25, p>0.05), or at the 24-hour time point (t_11_=0.87, p>0.05). n=12.

**Figure 1 – Supplement 3.**
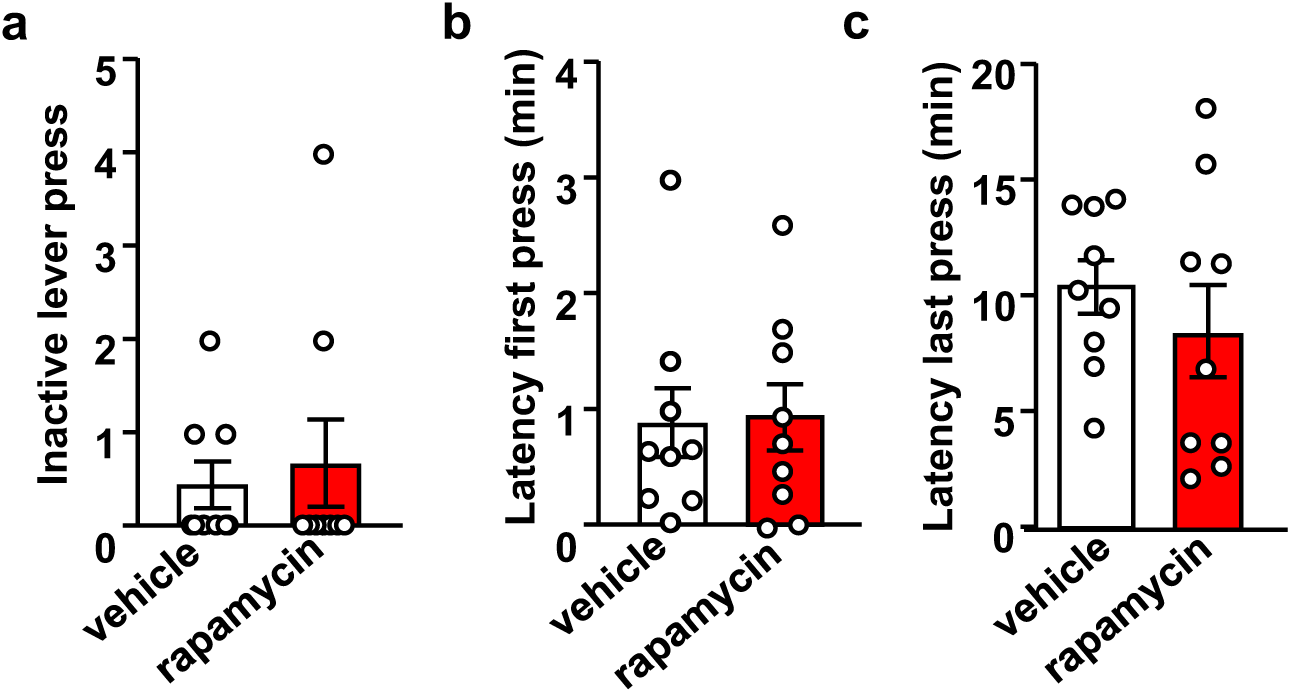
Inhibition of mTORC1 in the OFC does not impact inactive lever press or the latency to the first and last active lever press. Rats were trained to self-administer 20% alcohol using an FR3 schedule. Vehicle (white) or rapamycin (50ng/μl, red) was infused into the OFC 3 hours before an operant self-administration session and active and inactive lever presses were recorded. Significance between treatment groups was determined using two-tailed paired t-tests. **(a)** number of inactive lever presses (t_8_=0.37, p>0.05), latency to first lever press **(b)** (t_8_=0.27, p>0.05), and latency to the last active lever press **(c)** (t_8_=0.89, p>0.05) did not differ between groups. n=9.

**Figure 1 – Supplement 4.**
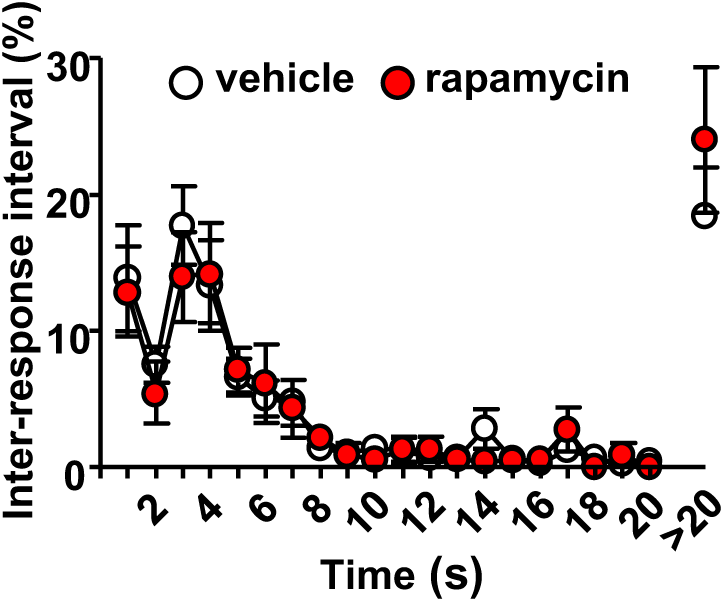
Intra-OFC infusion of rapamycin does not alter locomotion. Rats were trained to self-administer 20% alcohol using a FR3 schedule. Vehicle (white) or rapamycin (50ng/μl, red) was infused into the OFC 3 hours before a 30 minutes extinction session, and inter-response intervals (IRIs) were measured. Relative number of IRIs expressed as % of total IRIs with all intervals equal or smaller of 20 seconds presented in 1 second bins. Interval greater than 20 seconds are also shown. Two-way ANOVA of IRIs across the session did not identify a significant main treatment effect (F_1,12_=0.1412, p=0.7137). n=9.

**Figure 2 – Supplement 1.**
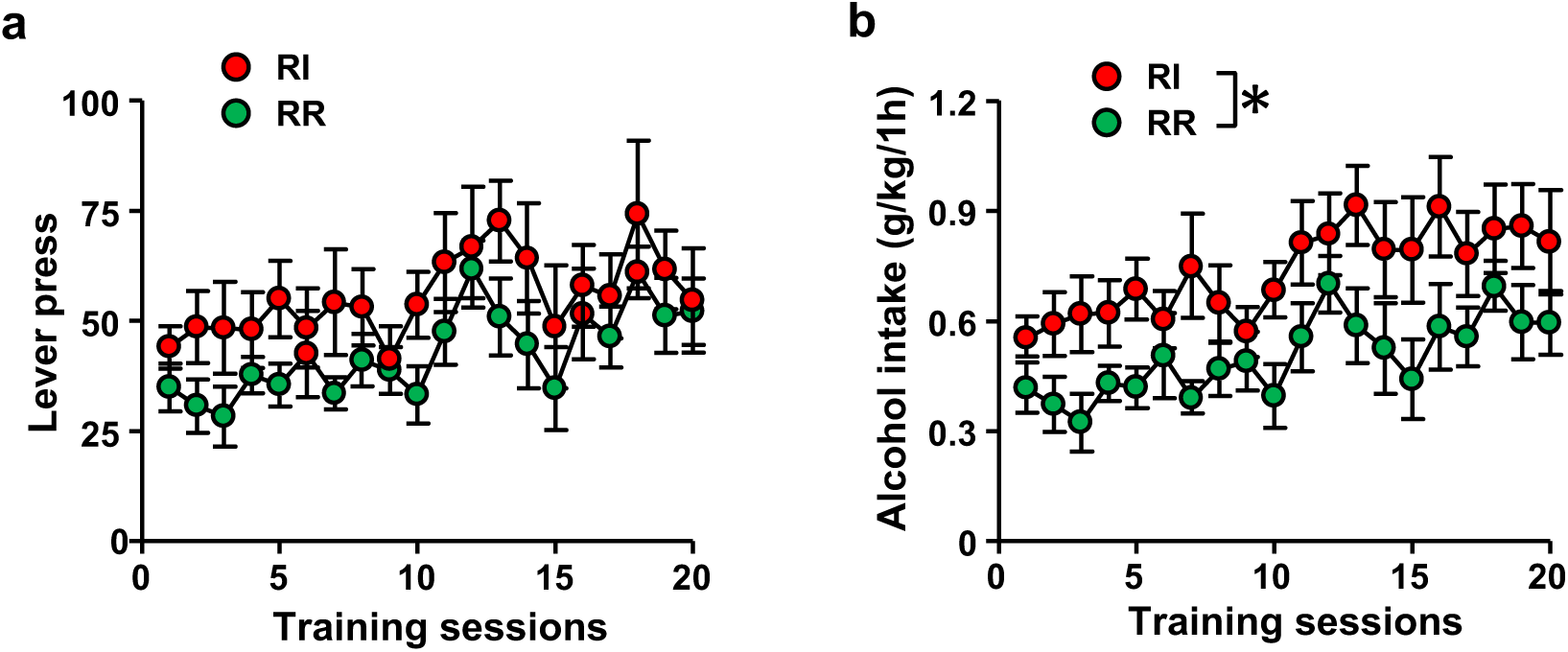
Number of lever presses and alcohol consumed in RI and RR-trained animals. Rats were trained to respond for alcohol in an operant self-administration paradigm under RI (red) or RR, (green) schedule of reinforcement. Significance between treatment groups was determined using RM ANOVA. **(a)** Training groups did not differ in the number of lever presses during training (F_1,20_=2.01, p>0.05), **(b)** RI-trained rats exhibited a small but significant increase of alcohol intake over the course of training (F_1,20_=6.65, p<0.05). *p<0.05. RR n=12. RI n=10.

**Figure 2 – Supplement 2.**
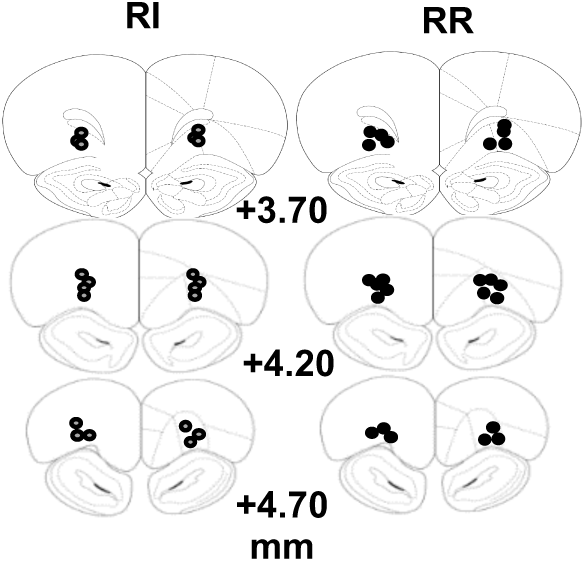
Schematic drawing of cannulae placement. Schematic illustration of coronal sections of the rat brain showing approximate bilateral placements of cannulae in the OFC according to Paxinos and Watson 2007.

**Figure 2 – Supplement 3.**
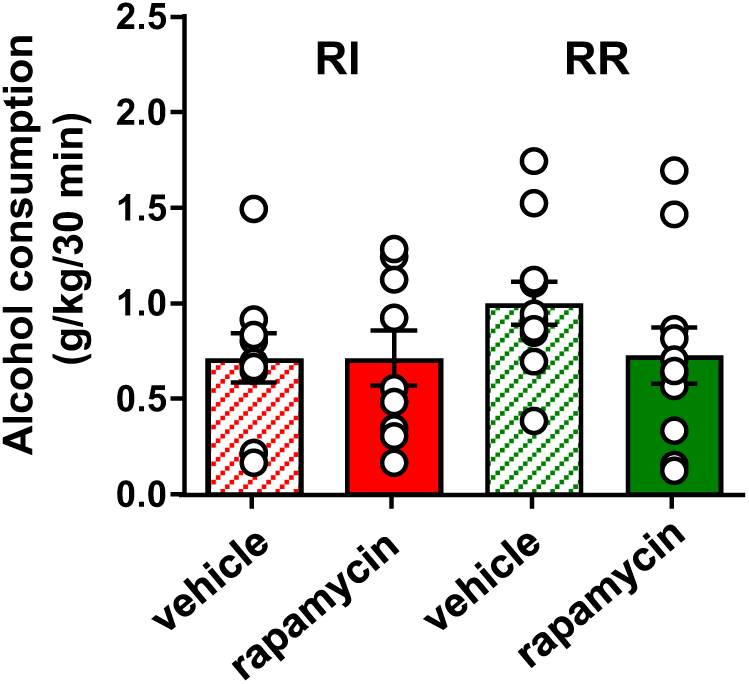
Inhibition of mTORC1 in the OFC does not alter voluntary alcohol intake prior to devaluation. Rats were trained to respond for alcohol in an operant self-administration paradigm under a random interval (RI, red) or random ration (RR, green) schedule of reinforcement. Vehicle (white) or rapamycin (50ng/μl, red) was infused into the OFC 3 hours prior to home cage access to alcohol, and alcohol intake was recorded at the end of the 30 minutes session. Significance between treatment groups was determined using RM ANOVA. The amount of alcohol consumed did not differ by treatment (F_1,30_=0.21, p>0.05) or training history (F_1,30_=1.48, p>0.05). *p<0.05. RR n=12. RI n=10.

**Figure 3 – Supplement 1.**
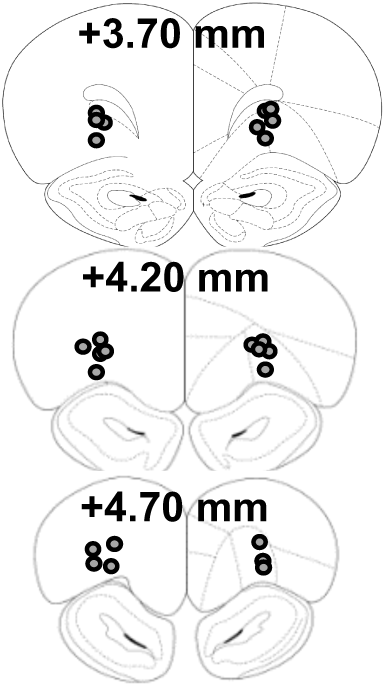
Schematic drawing of cannulae placement. Schematic illustration of coronal sections of the rat brain showing approximate bilateral placements of cannulae in the OFC according to Paxinos and Watson 2007.

**Figure 3 – Supplement 2.**
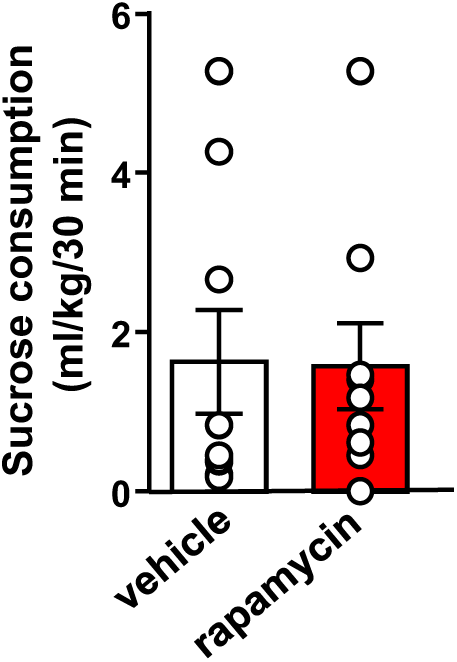
Inhibition of mTORC1 in the OFC does not alter voluntary sucrose intake prior to devaluation. Rats were trained to respond for sucrose in an operant self-administration paradigm under RI schedule of reinforcement. Vehicle (white) or rapamycin (50ng/μl, red) was infused into the OFC 3 hours prior to home cage access to sucrose, and sucrose intake was recorded at the end of the 30 minutes session. Significance between treatment groups was determined using RM ANOVA. The amount of sucrose consumed did not differ by treatment (F_1,30_=0.06, p>0.05). n=8.

**Figure 4 – Supplement 1.**
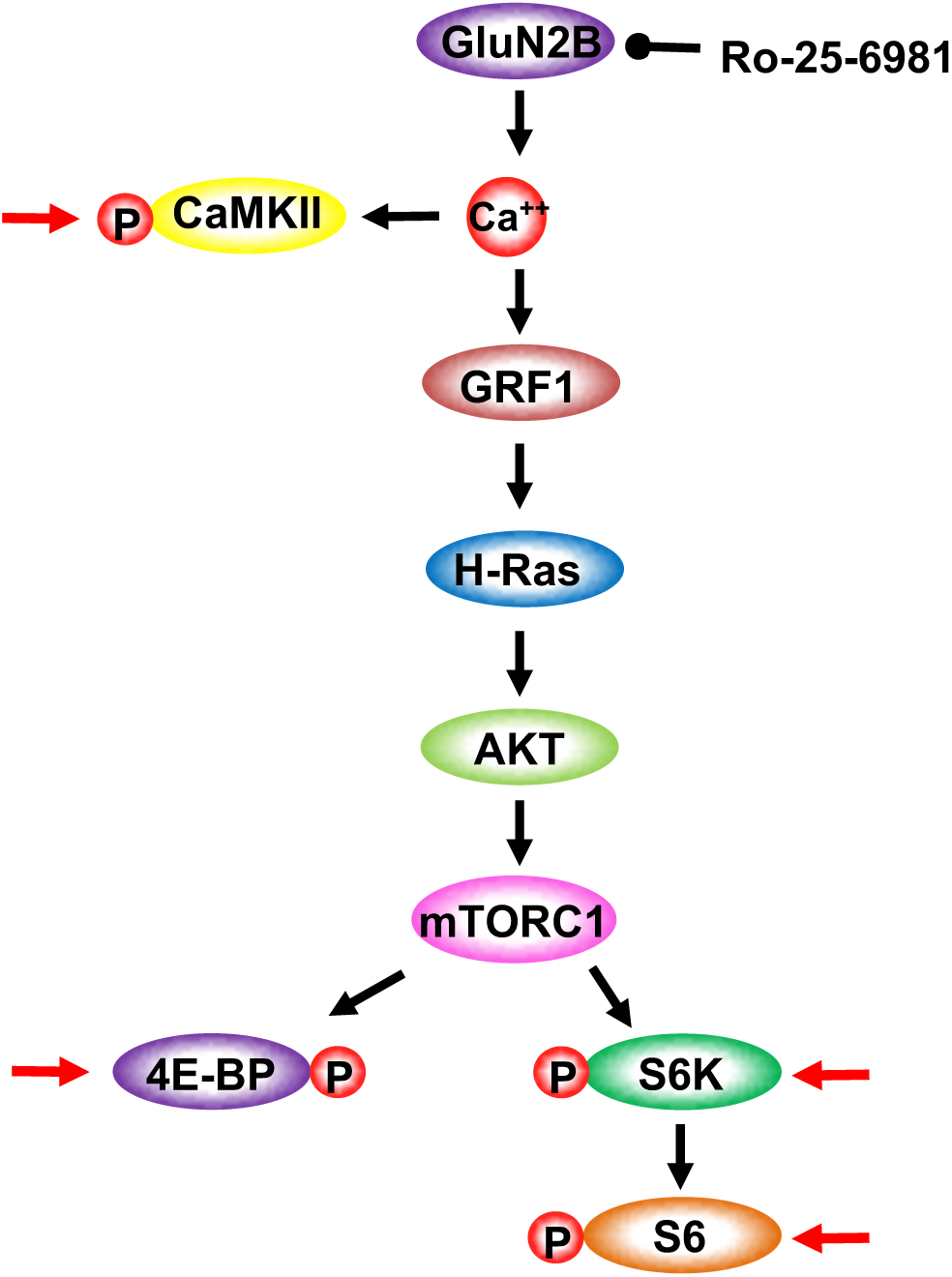
GluN2B-dependent activation of mTORC1 signaling. Model depicts the signaling cascade associated with Figure 4. GluN2B activation results in calcium release which in turn activates CaMKII and GRF1. CamKII activation is measured by phosphorylation (P) was used as a positive control. Calium binding to GRF1 activates H-Ras which activates AKT that in turn activates mTORC1. mTORC1 activation is measured by the phosphorylation of 4E-BP, S6K and S6. Red arrows depicts the specific proteins whose phosphorylation levels were tested.

**Figure 4 – Supplement 2.**
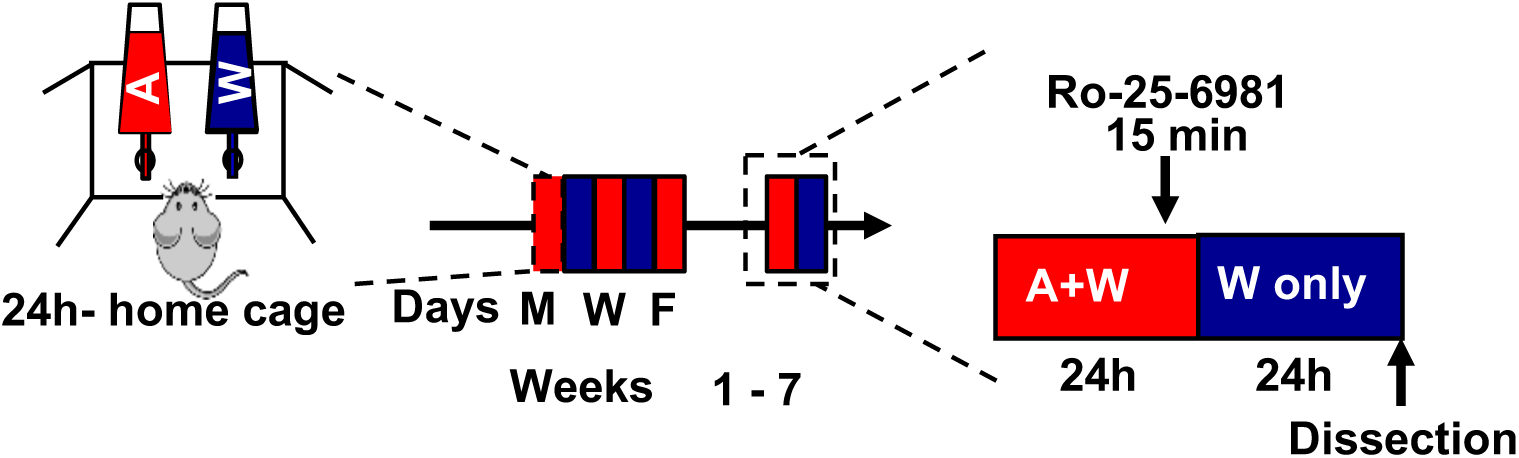
Time line of experiments depicted in Figure 4c-h. Rats underwent 7-weeks of IA-20%2BC paradigm or drank water only for the same duration of time. Ro-25-6981 (53) (10μg/ul) or vehicle was infused into the OFC infusion 15 minutes before the end of the last 24 hours alcohol drinking session, and mTORC1 activation was measured 24 hours later.

**Figure 5 – Supplement 1.**
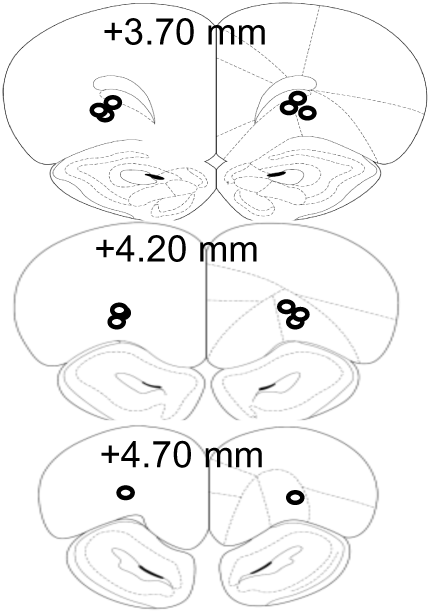
Schematic drawing of cannulae placement. Schematic illustration of coronal sections of the rat brain showing approximate bilateral placements of cannulae in the OFC according to Paxinos and Watson 2007.

**Figure 5 – Supplement 2.**
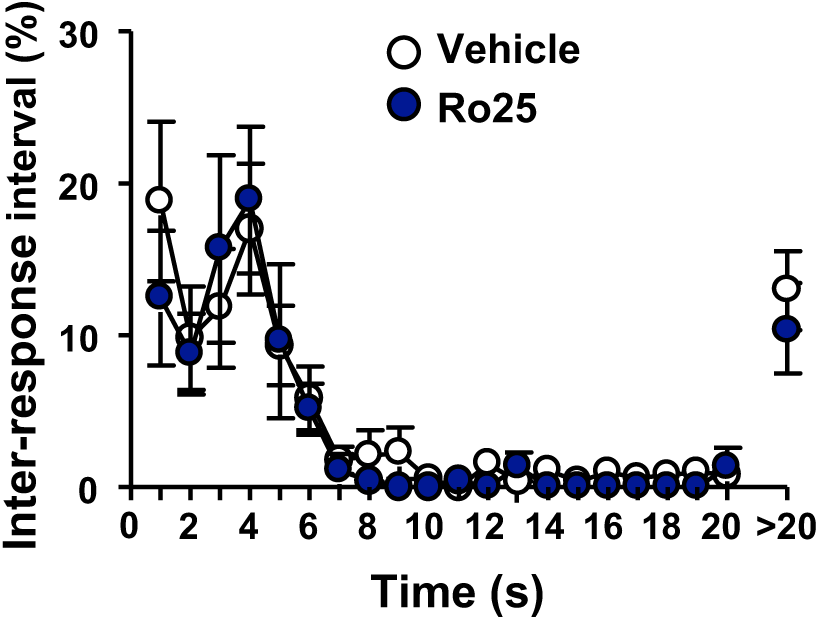
Inhibition of GluN2B in the OFC does not alter locomotion. Vehicle (white) or Ro25-6981 (10μg/μl, blue) was infused in the OFC 15 minutes before a single 30 minutes extinction session and inter-response intervals (IRIs) were measured. Mean +/- SEM of the relative number of IRIs expressed as % of total IRIs with all intervals equal or smaller of 20 seconds presented in 1 second bins. Intervals greater than 20 seconds are also shown. Two way RM ANOVA of IRIs across the session did not identify a significant main treatment effect (F_1,12_=0.81, p=0.3858). n=7.

**Figure 5 – Supplement 3.**
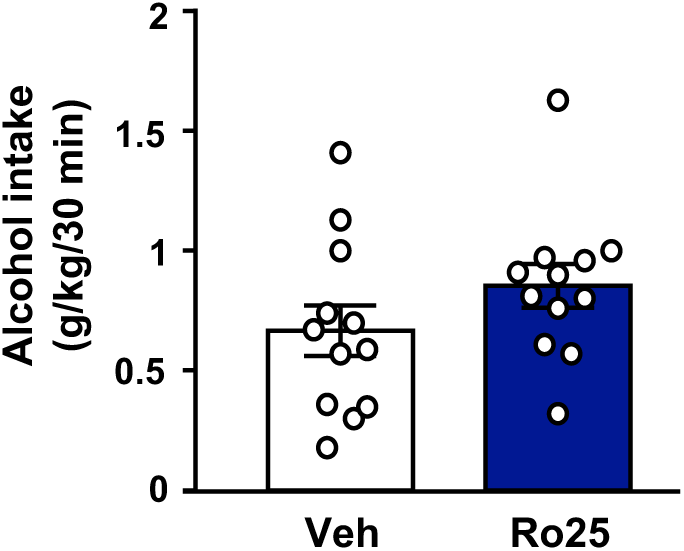
Inhibition of GluN2B in the OFC does not alter voluntary alcohol intake prior to devaluation. Rats were trained to respond for alcohol in an operant self-administration paradigm under a random interval (RI) schedule of reinforcement. Vehicle (white) or Ro25-6981 (5μg/μl, blue) was infused into the OFC 15 minutes prior to home cage access to alcohol, and alcohol intake was recorded at the end of the 30 minutes session. Significance between treatment groups was determined using RM ANOVA. The amount of alcohol consumed did not differ by treatment (t_5_=2.80, p>0.05). n=12.

## REFERENCES

1. Laguesse S, Morisot N, Phamluong K, Ron D (2017): Region specific activation of the AKT and mTORC1 pathway in response to excessive alcohol intake in rodents. Addict Biol. 22:1856–1869.

2. Lipton JO, Sahin M (2014): The neurology of mTOR. Neuron. 84:275–291.

3. Saxton RA, Sabatini DM (2017): mTOR Signaling in Growth, Metabolism, and Disease. Cell. 169:361–371.

4. Buffington SA, Huang W, Costa-Mattioli M (2014): Translational control in synaptic plasticity and cognitive dysfunction. Annu Rev Neurosci. 37:17–38.

5. Santini E, Huynh TN, Klann E (2014): Mechanisms of translation control underlying long-lasting synaptic plasticity and the consolidation of long-term memory. Prog Mol Biol Transl Sci. 122:131–167.

6. Hoeffer CA, Klann E (2010): mTOR signaling: at the crossroads of plasticity, memory and disease. Trends Neurosci. 33:67–75.

7. Neasta J, Barak S, Hamida SB, Ron D (2014): mTOR complex 1: a key player in neuroadaptations induced by drugs of abuse. J Neurochem. 130:172–184.

8. Neasta J, Ben Hamida S, Yowell Q, Carnicella S, Ron D (2010): Role for mammalian target of rapamycin complex 1 signaling in neuroadaptations underlying alcohol-related disorders. Proc Natl Acad Sci U S A. 107:20093–20098.

9. Beckley JT, Laguesse S, Phamluong K, Morisot N, Wegner SA, Ron D (2016): The First Alcohol Drink Triggers mTORC1-Dependent Synaptic Plasticity in Nucleus Accumbens Dopamine D1 Receptor Neurons. J Neurosci. 36:701–713.

10. Ben Hamida S, Laguesse S, Morisot N, Park JH, Phuamluong K, Berger AL, et al. (2019): Mammalian target of rapamycin complex 1 and its downstream effector collapsin response mediator protein-2 drive reinstatement of alcohol reward seeking. Addict Biol. 24:908–920.

11. Ben Hamida S, Laguesse S, Morisot N, Park JH, Phuamluong K, Berger AL, et al. (2018): Mammalian target of rapamycin complex 1 and its downstream effector collapsin response mediator protein-2 drive reinstatement of alcohol reward seeking. Addict Biol. 24(5):908–920.

12. Liu F, Laguesse S, Legastelois R, Morisot N, Ben Hamida S, Ron D (2017): mTORC1-dependent translation of collapsin response mediator protein-2 drives neuroadaptations underlying excessive alcohol-drinking behaviors. Mol Psychiatry. 22:89–101.

13. Wallis JD (2007): Orbitofrontal cortex and its contribution to decision-making. Annu Rev Neurosci. 30:31–56.

14. Meyer HC, Bucci DJ (2016): Imbalanced Activity in the Orbitofrontal Cortex and Nucleus Accumbens Impairs Behavioral Inhibition. Curr Biol. 26:2834–2839.

15. Baltz ET, Yalcinbas EA, Renteria R, Gremel CM (2018): Orbital frontal cortex updates state-induced value change for decision-making. Elife. 7. doi: 10.7554/eLife.35988.

16. Gremel CM, Costa RM (2013): Orbitofrontal and striatal circuits dynamically encode the shift between goal-directed and habitual actions. Nat Commun. 4:2264.

17. Ostlund SB, Balleine BW (2007): Orbitofrontal cortex mediates outcome encoding in Pavlovian but not instrumental conditioning. J Neurosci. 27:4819–4825.

18. Gourley SL, Olevska A, Zimmermann KS, Ressler KJ, Dileone RJ, Taylor JR (2013): The orbitofrontal cortex regulates outcome-based decision-making via the lateral striatum. Eur J Neurosci. 38:2382–2388.

19. Fiuzat EC, Rhodes SE, Murray EA (2017): The Role of Orbitofrontal-Amygdala Interactions in Updating Action-Outcome Valuations in Macaques. J Neurosci. 37:2463–2470.

20. Bradfield LA, Dezfouli A, van Holstein M, Chieng B, Balleine BW (2015): Medial Orbitofrontal Cortex Mediates Outcome Retrieval in Partially Observable Task Situations. Neuron. 88:1268–1280.

21. Gremel CM, Chancey JH, Atwood BK, Luo G, Neve R, Ramakrishnan C, et al. (2016): Endocannabinoid Modulation of Orbitostriatal Circuits Gates Habit Formation. Neuron. 90:1312–1324.

22. Ahmari SE, Spellman T, Douglass NL, Kheirbek MA, Simpson HB, Deisseroth K, et al. (2013): Repeated cortico-striatal stimulation generates persistent OCD-like behavior. Science. 340:1234–1239.

23. Pascoli V, Hiver A, Van Zessen R, Loureiro M, Achargui R, Harada M, et al. (2018): Stochastic synaptic plasticity underlying compulsion in a model of addiction. Nature. 564:366–371.

24. Schoenbaum G, Shaham Y (2008): The role of orbitofrontal cortex in drug addiction: a review of preclinical studies. Biol Psychiatry. 63:256–262.

25. Everitt BJ, Hutcheson DM, Ersche KD, Pelloux Y, Dalley JW, Robbins TW (2007): The orbital prefrontal cortex and drug addiction in laboratory animals and humans. Ann N Y Acad Sci. 1121:576–597.

26. Luscher C (2016): The Emergence of a Circuit Model for Addiction. Annu Rev Neurosci. 39:257–276.

27. McGuier NS, Padula AE, Lopez MF, Woodward JJ, Mulholland PJ (2015): Withdrawal from chronic intermittent alcohol exposure increases dendritic spine density in the lateral orbitofrontal cortex of mice. Alcohol. 49:21–27.

28. Nimitvilai S, Lopez MF, Mulholland PJ, Woodward JJ (2016): Chronic Intermittent Ethanol Exposure Enhances the Excitability and Synaptic Plasticity of Lateral Orbitofrontal Cortex Neurons and Induces a Tolerance to the Acute Inhibitory Actions of Ethanol. Neuropsychopharmacology. 41:1112–1127.

29. den Hartog C, Zamudio-Bulcock P, Nimitvilai S, Gilstrap M, Eaton B, Fedarovich H, et al. (2016): Inactivation of the lateral orbitofrontal cortex increases drinking in ethanol-dependent but not non-dependent mice. Neuropharmacology. 107:451–459.

30. Nimitvilai S, Uys JD, Woodward JJ, Randall PK, Ball LE, Williams RW, et al. (2017): Orbitofrontal Neuroadaptations and Cross-Species Synaptic Biomarkers in Heavy-Drinking Macaques. J Neurosci. 37:3646–3660.

31. Pfefferbaum A, Sullivan EV (2005): Disruption of brain white matter microstructure by excessive intracellular and extracellular fluid in alcoholism: evidence from diffusion tensor imaging. Neuropsychopharmacology. 30:423–432.

32. Miguel-Hidalgo JJ, Overholser JC, Meltzer HY, Stockmeier CA, Rajkowska G (2006): Reduced glial and neuronal packing density in the orbitofrontal cortex in alcohol dependence and its relationship with suicide and duration of alcohol dependence. Alcohol Clin Exp Res. 30:1845–1855.

33. Dom G, Sabbe B, Hulstijn W, van den Brink W (2005): Substance use disorders and the orbitofrontal cortex: systematic review of behavioural decision-making and neuroimaging studies. Br J Psychiatry. 187:209–220.

34. Claus ED, Ewing SWF, Filbey FM, Sabbineni A, Hutchison KE (2011): Identifying neurobiological phenotypes associated with alcohol use disorder severity. Neuropsychopharmacology. 36:2086–2096.

35. Filbey FM, Claus E, Audette AR, Niculescu M, Banich MT, Tanabe J, et al. (2008): Exposure to the taste of alcohol elicits activation of the mesocorticolimbic neurocircuitry. Neuropsychopharmacology. 33:1391–1401.

36. Laguesse S, Morisot N, Shin JH, Liu F, Adrover MF, Sakhai SA, et al. (2017): Prosapip1-Dependent Synaptic Adaptations in the Nucleus Accumbens Drive Alcohol Intake, Seeking, and Reward. Neuron. 96:145–159 e148.

37. Jeanblanc J, He DY, Carnicella S, Kharazia V, Janak PH, Ron D (2009): Endogenous BDNF in the dorsolateral striatum gates alcohol drinking. J Neurosci. 29:13494–13502.

38. Corbit LH, Nie H, Janak PH (2012): Habitual alcohol seeking: time course and the contribution of subregions of the dorsal striatum. Biol Psychiatry. 72:389–395.

39. Renteria R, Baltz ET, Gremel CM (2018): Chronic alcohol exposure disrupts top-down control over basal ganglia action selection to produce habits. Nat Commun. 9:211.

40. Sutton G, Chandler LJ (2002): Activity-dependent NMDA receptor-mediated activation of protein kinase B/Akt in cortical neuronal cultures. J Neurochem. 82:1097–1105.

41. Barak S, Liu F, Ben Hamida S, Yowell QV, Neasta J, Kharazia V, et al. (2013): Disruption of alcohol-related memories by mTORC1 inhibition prevents relapse. Nat Neurosci. 16:1111–1117.

42. Carnicella S, Kharazia V, Jeanblanc J, Janak PH, Ron D (2008): GDNF is a fast-acting potent inhibitor of alcohol consumption and relapse. Proc Natl Acad Sci U S A. 105:8114–8119.

43. Corbit LH, Janak PH (2016): Habitual Alcohol Seeking: Neural Bases and Possible Relations to Alcohol Use Disorders. Alcohol Clin Exp Res. 40:1380–1389.

44. Seif T, Chang SJ, Simms JA, Gibb SL, Dadgar J, Chen BT, et al. (2013): Cortical activation of accumbens hyperpolarization-active NMDARs mediates aversion-resistant alcohol intake. Nat Neurosci. 16:1094–1100.

45. Pascoli V, Terrier J, Hiver A, Luscher C (2015): Sufficiency of Mesolimbic Dopamine Neuron Stimulation for the Progression to Addiction. Neuron. 88:1054–1066.

46. Rossi MA, Yin HH (2012): Methods for studying habitual behavior in mice. Curr Protoc Neurosci. Chapter 8:Unit 8 29.

47. Schoenbaum G, Setlow B, Saddoris MP, Gallagher M (2003): Encoding predicted outcome and acquired value in orbitofrontal cortex during cue sampling depends upon input from basolateral amygdala. Neuron. 39:855–867.

48. Lichtenberg NT, Pennington ZT, Holley SM, Greenfield VY, Cepeda C, Levine MS, et al. (2017): Basolateral Amygdala to Orbitofrontal Cortex Projections Enable Cue-Triggered Reward Expectations. J Neurosci. 37:8374–8384.

49. Hermann D, Weber-Fahr W, Sartorius A, Hoerst M, Frischknecht U, Tunc-Skarka N, et al. (2012): Translational magnetic resonance spectroscopy reveals excessive central glutamate levels during alcohol withdrawal in humans and rats. Biol Psychiatry. 71:1015–1021.

50. Morisot N, Ron D (2017): Alcohol-dependent molecular adaptations of the NMDA receptor system. Genes Brain Behav. 16:139–148.

51. Krapivinsky G, Krapivinsky L, Manasian Y, Ivanov A, Tyzio R, Pellegrino C, et al. (2003): The NMDA receptor is coupled to the ERK pathway by a direct interaction between NR2B and RasGRF1. Neuron. 40:775–784.

52. Manning BD, Toker A (2017): AKT/PKB Signaling: Navigating the Network. Cell. 169:381–405.

53. Fischer G, Mutel V, Trube G, Malherbe P, Kew JN, Mohacsi E, et al. (1997): Ro 25-6981, a highly potent and selective blocker of N-methyl-D-aspartate receptors containing the NR2B subunit. Characterization in vitro. J Pharmacol Exp Ther. 283:1285–1292.

54. Schoenbaum G, Chiba AA, Gallagher M (1998): Orbitofrontal cortex and basolateral amygdala encode expected outcomes during learning. Nat Neurosci. 1:155–159.

55. O’Doherty JP, Deichmann R, Critchley HD, Dolan RJ (2002): Neural responses during anticipation of a primary taste reward. Neuron. 33:815–826.

56. Belin D, Belin-Rauscent A, Murray JE, Everitt BJ (2013): Addiction: failure of control over maladaptive incentive habits. Curr Opin Neurobiol. 23:564–572.

57. Everitt BJ, Robbins TW (2005): Neural systems of reinforcement for drug addiction: from actions to habits to compulsion. Nat Neurosci. 8:1481–1489.

58. Fettes P, Schulze L, Downar J (2017): Cortico-Striatal-Thalamic Loop Circuits of the Orbitofrontal Cortex: Promising Therapeutic Targets in Psychiatric Illness. Front Syst Neurosci. 11:25.

59. Izquierdo A (2017): Functional Heterogeneity within Rat Orbitofrontal Cortex in Reward Learning and Decision Making. J Neurosci. 37:10529–10540.

60. Noonan MP, Walton ME, Behrens TE, Sallet J, Buckley MJ, Rushworth MF (2010): Separate value comparison and learning mechanisms in macaque medial and lateral orbitofrontal cortex. Proc Natl Acad Sci U S A. 107:20547–20552.

61. Smith RJ, Laiks LS (2018): Behavioral and neural mechanisms underlying habitual and compulsive drug seeking. Prog Neuropsychopharmacol Biol Psychiatry. 87:11–21.

62. Warnault V, Darcq E, Morisot N, Phamluong K, Wilbrecht L, Massa SM, et al. (2016): The BDNF Valine 68 to Methionine Polymorphism Increases Compulsive Alcohol Drinking in Mice That Is Reversed by Tropomyosin Receptor Kinase B Activation. Biol Psychiatry. 79:463–473.

63. Augier E, Barbier E, Dulman RS, Licheri V, Augier G, Domi E, et al. (2018): A molecular mechanism for choosing alcohol over an alternative reward. Science. 360:1321–1326.

64. Enoch MA, Goldman D (2002): Problem drinking and alcoholism: diagnosis and treatment. Am Fam Physician. 65:441–448.

65. Carnicella S, Ron D, Barak S (2014): Intermittent ethanol access schedule in rats as a preclinical model of alcohol abuse. Alcohol. 48:243–252.

66. NIAAA (2004): NIAAA council approves definition of binge drinking. *NIAAA Newsletter, Winter 2004, Number 3*. Bethesda, MD: Office of Research Translation and Communications, NIAAA, NIH, DHHS, pp 3.

67. Griffin WC, 3rd (2014): Alcohol dependence and free-choice drinking in mice. Alcohol. 48:287–293.

68. WHO (2014): Global status report on alcohol and health. World Health Organization.

69. Trepanier CH, Jackson MF, MacDonald JF (2012): Regulation of NMDA receptors by the tyrosine kinase Fyn. FEBS J. 279:12–19.

70. Yoon YJ, Wu B, Buxbaum AR, Das S, Tsai A, English BP, et al. (2016): Glutamate-induced RNA localization and translation in neurons. Proc Natl Acad Sci U S A. 113:E6877–E6886.

71. DePoy LM, Zimmermann KS, Marvar PJ, Gourley SL (2017): Induction and Blockade of Adolescent Cocaine-Induced Habits. Biol Psychiatry. 81:595–605.

72. Brigman JL, Daut RA, Wright T, Gunduz-Cinar O, Graybeal C, Davis MI, et al. (2013): GluN2B in corticostriatal circuits governs choice learning and choice shifting. Nat Neurosci. 16:1101–1110.

73. Puighermanal E, Marsicano G, Busquets-Garcia A, Lutz B, Maldonado R, Ozaita A (2009): Cannabinoid modulation of hippocampal long-term memory is mediated by mTOR signaling. Nat Neurosci. 12:1152–1158.

74. Gourley SL, Zimmermann KS, Allen AG, Taylor JR (2016): The Medial Orbitofrontal Cortex Regulates Sensitivity to Outcome Value. J Neurosci. 36:4600–4613.

75. Zimmermann KS, Yamin JA, Rainnie DG, Ressler KJ, Gourley SL (2017): Connections of the Mouse Orbitofrontal Cortex and Regulation of Goal-Directed Action Selection by Brain-Derived Neurotrophic Factor. Biol Psychiatry. 81:366–377.

76. Ron D, Barak S (2016): Molecular mechanisms underlying alcohol-drinking behaviours. Nat Rev Neurosci. 17:576–591.

